# Mathematical modeling with single-cell sequencing data

**DOI:** 10.1101/710640

**Authors:** Heyrim Cho, Russell C. Rockne

## Abstract

Single-cell sequencing technologies have revolutionized molecular and cellular biology and stimulated the development of computational tools to analyze the data generated from these technology platforms. However, despite the recent explosion of computational analysis tools, relatively few mathematical models have been developed to utilize these data. Here we compare and contrast two approaches for building mathematical models of cell state-transitions with single-cell RNA-sequencing data with hematopoeisis as a model system; by solving partial differential equations on a graph representing discrete cell state relationships, and by solving the equations on a continuous cell state-space. We demonstrate how to calibrate model parameters from single or multiple time-point single-cell sequencing data, and examine the effects of data processing algorithms on the model calibration and predictions. As an application of our approach, we demonstrate how the calibrated models may be used to mathematically perturb normal hematopoeisis to simulate, predict, and study the emergence of novel cell types during the pathogenesis of acute myeloid leukemia. The mathematical modeling framework we present is general and can be applied to study cell state-transitions in any single-cell genome sequencing dataset.

**Author summary:** Here we compare and contrast graph- and continuum-based approaches for constructing mathematical models of cell state-transitions using single-cell RNA-sequencing data. Using two publicly available datasets, we demonstrate how to calibrate mathematical models of hematopoeisis and how to use the models to predict dynamics of acute myeloid leukemia pathogenesis by mathematically perturbing the process of cellular proliferation and differentiation. We apply these modeling approaches to study the effects of perturbing individual or sets of genes in subsets of cells, or by modeling the dynamics of cell state-transitions directly in a reduced dimensional space. We examine the effects of different graph abstraction and trajectory inference algorithms on calibrating the models and the subsequent model predictions. We conclude that both the graph- and continuum-based modeling approaches can be equally well calibrated to data and discuss situations in which one method may be preferable over the other. This work presents a general mathematical modeling framework, applicable to any single-cell sequencing dataset where cell state-transitions are of interest.

## Introduction

The ability to apply genome sequencing methods to single cells has revolutionized biology [1]. Technologies enabling single-cell sequencing are advancing rapidly, with datasets as large as hundreds of thousands of cells [2]. RNA-sequencing is currently the most prevalent form of single-cell genomic analysis (scRNA-seq) [1]. The sequencing of RNA at the cellular level enables the interrogation of gene transcription, which is used as a high-dimensional fingerprint which characterizes the identity of the cell. For this reason, scRNA-seq has been used as a tool to study cell identity and state-transitions at the cellular level.

The most frequently studied cell state-transition is cellular differentiation; the process of a cell and its progeny to perform specialized tasks through transformation from a less differentiated stem-like state to a more differentiated state. ScRNA-seq is used to identify cells in various states of differentiation primarily through one or both of two primary methods: 1) clustering of cells with similar features [3], or 2) though trajectory inference (TI) [4]. Clustering analysis relies on the definition of a similarity metric, and may rely on a pre-defined number of clusters (ex. k-means), or may use optimization methods to identify clusters (ex. Leiden). There are a wide variety of clustering methods and similarity metrics to choose from, which may give drastically different results [5]. Similarly, trajectory inference methods may use pre-defined relationships between cells or may use optimization methods to identify these relationships to construct graphs which are then used to infer paths, or trajectories, between cell states. In addition, various approaches aim to characterize the cell fate landscape, for instance, by a parameterized landscape based on bifurcation analysis [6, 7], by using a measure of entropy of cell states: SCENT [8] and scEpath [9], or by mapping cells to a landscape on optimized parameters: HopLand [10] and Topslam [11].

A significant limitation of these approaches is if the graph structure and underlying relationships between the cells is unknown. As shown in a comprehensive review of trajectory inference methods by Saelens et al, most TI algorithms have difficulty inferring even simple graphs which may include cycles or disconnected subgraphs [4]. We suggest that a hypothesis-driven and mathematical approach to the analysis of scRNA-seq data to study cell state transitions is warranted.

Moreover, single cell genomic sequencing suffers from a number of challenges in analysis. Beyond the several choices to be made for even simple analyses such as clustering or visualization, the data may be sparse and incomplete. Gene “drop outs” and background signal (noise) can complicate differential expression and clustering analyses. For this reason, analysis of these data have remained fairly superficial despite the wealth of information contained in these high-dimensional datasets. Moreover, results obtained from analysis of single cell sequencing datasets may be very sensitive to choices in the method of analysis and parameters in algorithms. To date, single-cell sequencing data have not been effectively leveraged as inputs into mathematical models.

Here we compare two approaches of mathematical modeling cell state-transitions with scRNA-seq data. Building on our prior work [12], we model cell differentiation as as a continuous process with reaction-diffusion-advection partial differential equations (PDE) solved on: 1) an abstracted graph and 2) a continuous space. We compare and contrast these two approaches with hematopoeisis as the model system.

This manuscript is structured as follows: first we present the PDE model on a graph and in continuous space, then we apply the model to two datasets. We examine the impact of various graph construction and trajectory inference methods on the geometry of the cell state space, and solve the model on these geometries. We then use the model the study the effects of perturbing 1) the graph structure 2) expression of select subsets of genes 3) and cell state transition dynamics by perturbing flow on the graph or by modifying the dynamics in the continuous space. We predict novel dynamics of leukemia pathogenesis by perturbing normal hematopoeisis and conclude with a comparative analysis of our approach and description of future directions for mathematical modeling with single cell genomic sequencing data. A summary of our workflow is shown in Fig 1.

**Fig 1.**
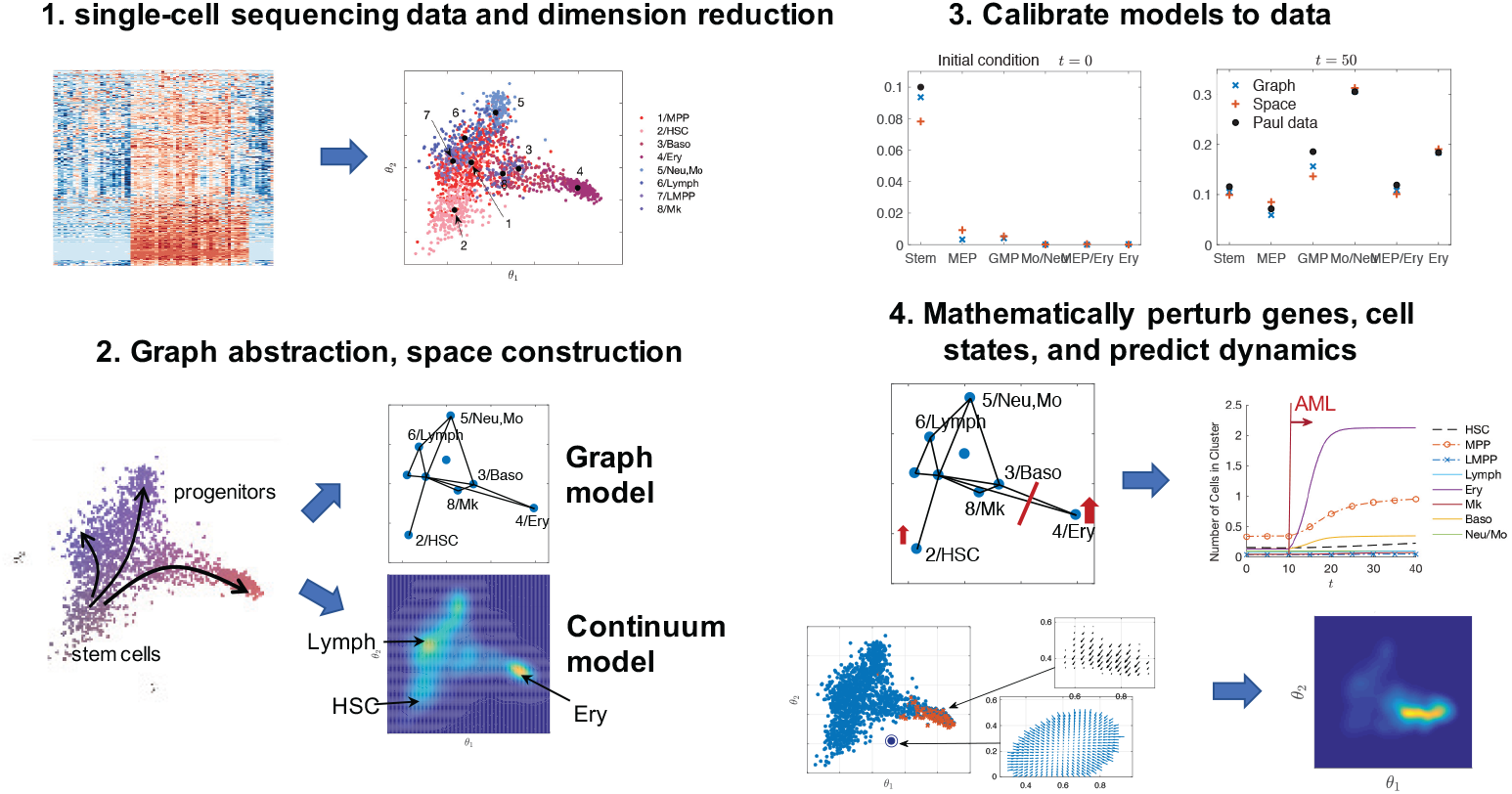
Step-by-step illustration of our modeling process. 1. Processed single-cell sequencing expression matrices are represented in a reduced dimension space through one of many dimension reduction techniques such as diffusion mapping, principle component analysis, t-SNE, or UMAP. 2. Temporal events (pseudotime trajectories) and cell clusters are inferred to construct the graph or continuum geometry of cell states. 3. From these representations, models are calibrated to the transport of cell distribution along the graph or in the cell state-space. 4. The models can then be utilized to perturb genes and cell states. The calibrated models predict cell state-transitions and the emergence of novel cell states.

## Materials and methods

### Modeling cell state-transitions in a continuous cell state-space

In this section we develop PDE models that describes the cell dynamics in the continuous phenotype space identified by dimension reduction techniques. For a given single cell genomic sequencing data 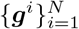, where *N* is the number of cell samples and ***g***^*i*^ ∈ ℝ^*G*^ is a *G*-dimensional vector of gene expression level of the *i*-th cell, the dimension reduction method can be written as an operator: 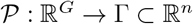 where the reduced dimensional space is truncated at the *n*-th dimension. Various dimension reduction techniques exist to potentially employ for the mapping 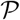, including principal component analysis, diffusion mapping, and t-distributed stochastic neighbor embedding. While different techniques provide different shapes and differentiation spaces, we choose diffusion mapping due to its ability to capture the non-linear structure of high-dimensional data, and to well reproduce global trajectory of data [13]. We denote the reduced space variable as *θ* = (*θ*_1_, *θ*_2_, …, *θ*_*n*_) *∈* Γ, and the *i*-th projected cell data in the reduced component space as 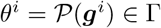.

### PDE model of cell state-transitions on a reduced component space

We first develop a cell state model that describes the dynamics of cell distribution *u*(*t, θ*) on the reduced component space Γ, where *θ* ϵ Γ is the variable that represents continuous cell state. Three highly distinctive dynamic regimes of the cell states are considered, that is, directed cell transition, random phenotypic instability, and birth-death process. Such model can be written as an advection-diffusion-reaction PDE that governs the cell distribution *u*(*t, θ*) as

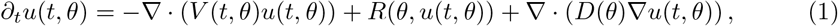

with zero Dirichlet boundary condition. The three terms in our equation that involve parameterized functions *V*, *R*, and *D*, represent cell differentiation, population growth, and phenotypic instability, respectively.

Let us first describe the advection term *V* ∈ ℝ^*n*^ that represents directed cell differentiation, where we divide it into two parts as *V* = **v**_1_ + **v**_2_. The first part models the attractor cell states of homeostasis. Assuming that the magnitude of phenotypic instability is with a magnitude *ν*, that is, *D*(*θ*) = *ν*, one can compute the advection term **v**_1_ as

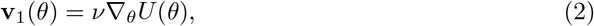

where *U* (*θ*) is the exponent of the homeostasis distribution in the exponential form *u*_*s*_(*θ*) = *e*^−*U*(*θ*)^. Here, *u*_*s*_(*θ*) can be regarded as the cell landscape that the hematopoiesis system desires to maintain. There are multiple methods to compute the cell landscape, so called quasi-potential [14], that focuses on relative stability of multiple attractors and models cell differentiation as transition between the attractor states. Here, we compute the cell landscape empirically by assuming that the obtained single-cell data is a representative subset of the entire hematopoiesis system, and by using density approximation methods. In particular, we use kernel density estimation [15] from the projected single-cell data *θ*^*i*^ Γ. The second advection term models an active cell differentiation. In particular, we consider the following form

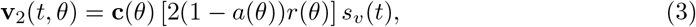

that is parameterized by the proliferation rate *r*(*θ*) and cell maturation [16]. **c**(*θ*) ϵ ℝ^*n*^ represents the direction and magnitude of differentiation on the phenotype space, and *a*(*θ*) is the proportion of cells that remains in cell type *θ*, while 1 − *a*(*θ*) cells further differentiate in to matured states. This is can be counted from symmetric and asymmetric stem cell division. In addition, we assume a signal parameter *s*_*v*_(*t*) that controls the active differentiation term.

The reaction term represents the growth rate that consist of proliferation and apoptosis. We consider the logistic growth term as following

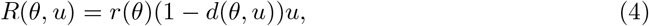

where *r*(*θ*) is the proliferation rate and *d*(*θ*) is the apoptosis term assuming a logistic growth as 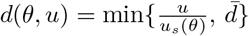, where 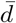 models the maximum magnitude of apoptosis rate. The second-order diffusion term represents the instability on the phenotypic landscape of the cells that should be taken account into when modeling the macroscopic cell density. We simply consider a constant term *D*(*θ*) = *ν*, and the value is estimated as *ν* ≈ *ε*^2^/4*δt* with *ε* as a single step size in the phenotype space and *δt* as the time elapsed between the steps. See S1 Appendix for the detail of the model.

### PDE model of cell state-transitions solved on a graph

Although the continuum-based multi-dimensional model provides a framework to study cell states, it is not always straightforward to map back locations in the space to novel or otherwise unknown cell types. Moreover, a central feature of contemporary analysis of scRNA-seq data is clustering and inferring relationships between clusters of known cell types [4]. Therefore, we develop a model that can describe cell state-transition dynamics on a graph that represents relationships between known cell types identified with clusters, extended from our previous work in [12]. An immediate advantage of this approach is that it is convenient to employ biological insights from well-known classical discrete cell states.

The continuum differentiation cell states are assumed to be on the graph obtained from the scRNA-seq data, for instance, using partition-based graph abstraction algorithm [17]. We project the graph on the reduced component space, and denote the nodes as 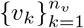 and the edges as *e*_*ij*_ connecting in the direction from the *i*-th to the *j*-th node. For convenience of notation, the edges are also denoted as 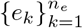 with the index mapping 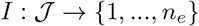 on the set of nontrivial edges (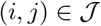. The end points in the direction of cell transition are 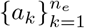 and 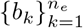, where 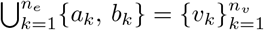.

The model follows the dynamics of the cell distribution on the graph, *u*(*x, t*), where *x* ∈ *e*_*k*_ is a one dimensional variable that parameterizes the differentiation continuum space location. We annotate the cell distribution on each edge *e*_*k*_ as *u*_*k*_(*x, t*) such that 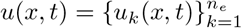, and model the cell density by an advection-diffusion-reaction equation [18] as

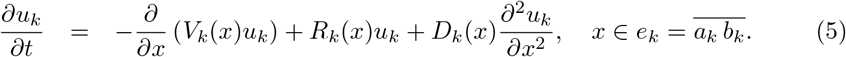

The three terms are similarly modeled as the multi-dimensional model in Eq (1), representing cell differentiation, population growth, and phenotypic instability. To summarize once more, the advection coefficient *V*_*k*_(*x*) models the cell differentiation and the transition between the nodes, that is, different cell types. We model the advection term in two parts as in the reduced component space model, *V*_*k*_(*x*) = *v*_*k*,1_(*x*) + *v*_*k*,2_(*x*), and compute them as

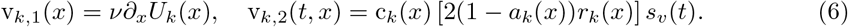

Here, 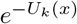 is the homeostasis cell distribution on the graph, *ν* is the magnitude of phenotypic instability, *r*_*k*_(*x*) is the proliferation rate, *a*_*k*_(*x*) is the self-renewal rate, and *s*_*v*_(*t*) is the signal parameter. The transition rate per unit time (e.g., day^*−*1^) or pseudotime can be taken into consideration in c_*k*_(*x*). Cell proliferation and apoptosis can be modeled by the reaction coefficient *R*_*k*_(*x*) as

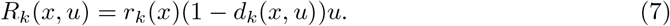

Finally, the diffusion term that represents phenotypic fluctuation is taken as *D*_*k*_(*x*) = *ν*. The magnitude of phenotypic fluctuation of the cell density is assumed as a random process subject to Brownian motion with magnitude *σ*, so that the diffusion term becomes *ν* = *σ*^2^*/*2 [18].

In addition to the governing equation on the edges, the boundary condition at the nodes are critical when describing the dynamics on the graph. The boundary condition on the cell fate PDE model can be classified into three types, the initial nodes that do not have inflow 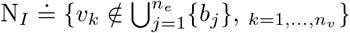, e.g., stem cells, the final nodes without outflow 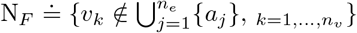, e.g., the most differentiated cells, and the intermediate nodes, 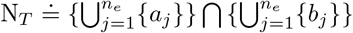. On the intermediate nodes *v*_*n*_ ∈ *N*_*T*_, mixed boundary condition is imposed for continuity of the density and flow as following.

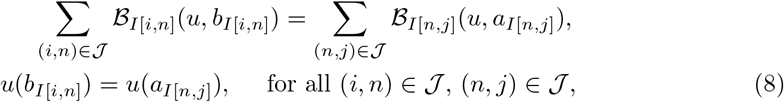

where 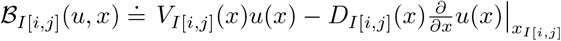, *b*_*I*[*i,n*]_ is the right end point of the edge between nodes *i* and *n*, and *a*_*I*[*n,j*]_ is the left end point of the edge between nodes *n* and *j*. The cell outflow boundary conditions on the final nodes, *v*_*n*_ ∈ N_*F*_, are imposed as reflecting boundary conditions

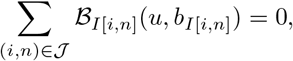

and *u*(*b*_*I*[*i,n*]_) = *u*(*b*_*I*[*j,n*]_) for all (*i, n*) and (*j, n*) in 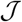, and similarly on the initial nodes *v*_*n*_ ∈ N_*I*_.

### Quantification of cell state-transition dynamics

Let us define some useful quantities to interpret model predictions in the continuous cell state-space and on a graph. The total number of cells from the cell distribution on either a graph or a continuous manifold can be computed as

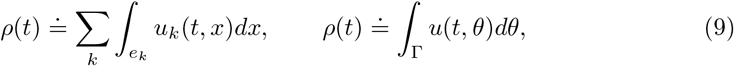

respectively. In addition, we compute the number of cells in specific cell type by assigning weight *w*_*k*_ that corresponds to cells in the *k*-th cluster as

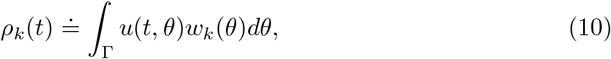

Σ_*k*_ *w*_*k*_(*θ*) = 1. In the graph model, we assign the cell states along the edge to be the cell type of the closest node, so that we can compute the number of the *k*-th node cell type as

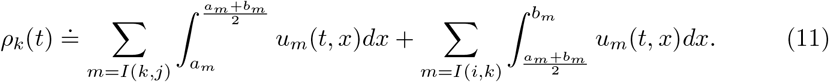

Although we can understand the continuum of cell states by mapping cells in intermediate states back to known discrete cell types, we also desire to interpret the new cell states along the manifold in their location without reference to the canonical cell identities. For such purpose, we characterize new cell states by identifying genes that are strongly correlated to a location in the space or moving in a direction toward a cell state. First, to characterize the cell state *θ*^∗^ in the reduced space, we use function 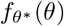 centered at *θ*^∗^ as 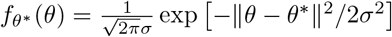, and compute the correlation between the function values and the *j*-th gene expression levels as

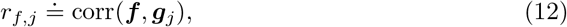

where ***f*** represents the vector of function evaluation at each single-cell data point *θ*^*i*^, that is, 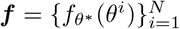, and 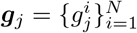. An alternate quantity to examine is the genes that are related to a certain direction in the reduced component space. For instance, the correlation between the *j*-th gene and the *k*-th reduced component 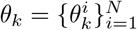 and to a certain vector 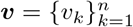 can be computed as

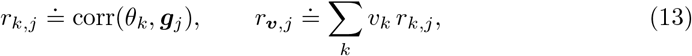

respectively. Regarding Eq (13) as global quantities, we can also compute the local correlation on the subdomain of the reduced space Ω_*d*_ by collecting the cell indices that lie within),the subdomain Γ_*d*_ = {*i* | *θ*^*i*^ ∈ Ω_*d*_}, that is, 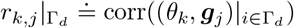 and 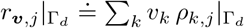.

## Results

In this section, we employ the framework developed in the previous section to the mouse hematopoiesis cell data from Nestorowa et al [19] and Paul et al [20]. We obtain the graph and space models of hematopoiesis cell state and focus on comparing the strengths and weaknesses of the two models.

### Comparing a continuous cell state-space with a cell state graph

The hematopoiesis single cell data from [19] and [20] projected on the first two diffusion component space are shown in Fig 2A, where distinct cell types, including lymphoid-primed multipotent progenitors (Lymph); common myeloid progenitors (CMP); megakaryocyte-erythroid progenitors (MEP); granulocyte-macrophage progenitors (GMP); erythrocytes (Ery); neutrophilic (Neu); monocyte (Mo); megakaryocytes (Mk); basophilic (Baso), classified in the original papers are illustrated with different colors. We truncate the diffusion component at two since the reduced two-dimensional space describes the dynamics of our interest, that is, from strong to weak stemness. The first two diffusion components *θ*_1_ and *θ*_2_ represent cell maturation in both data sets. In Nestorowa data, the first diffusion component separates the stem cells to myeloid lineages, particularly MEP cells and the second diffusion component describes GMP cells and the lymphoid lineages. In Paul data, the first and second reduced component represents MEP and GMP lineage, respectively. We remark that the cells that are the most stemlike in Paul data are common myeloid progenitors, that is more matured than the ones in Nestorowa data, that includes the long-term and short-term HSCs. In addition to the single-cell data, the figure also presents the abstracted graphs obtained using partition-based graph abstraction [17]. Further refinements of the graph will eventually become the full single-cell data, where each single-cell being counted as distinct cell states, and it depicts the hierarchy of cell states (see S7 Fig).

**Fig 2.**
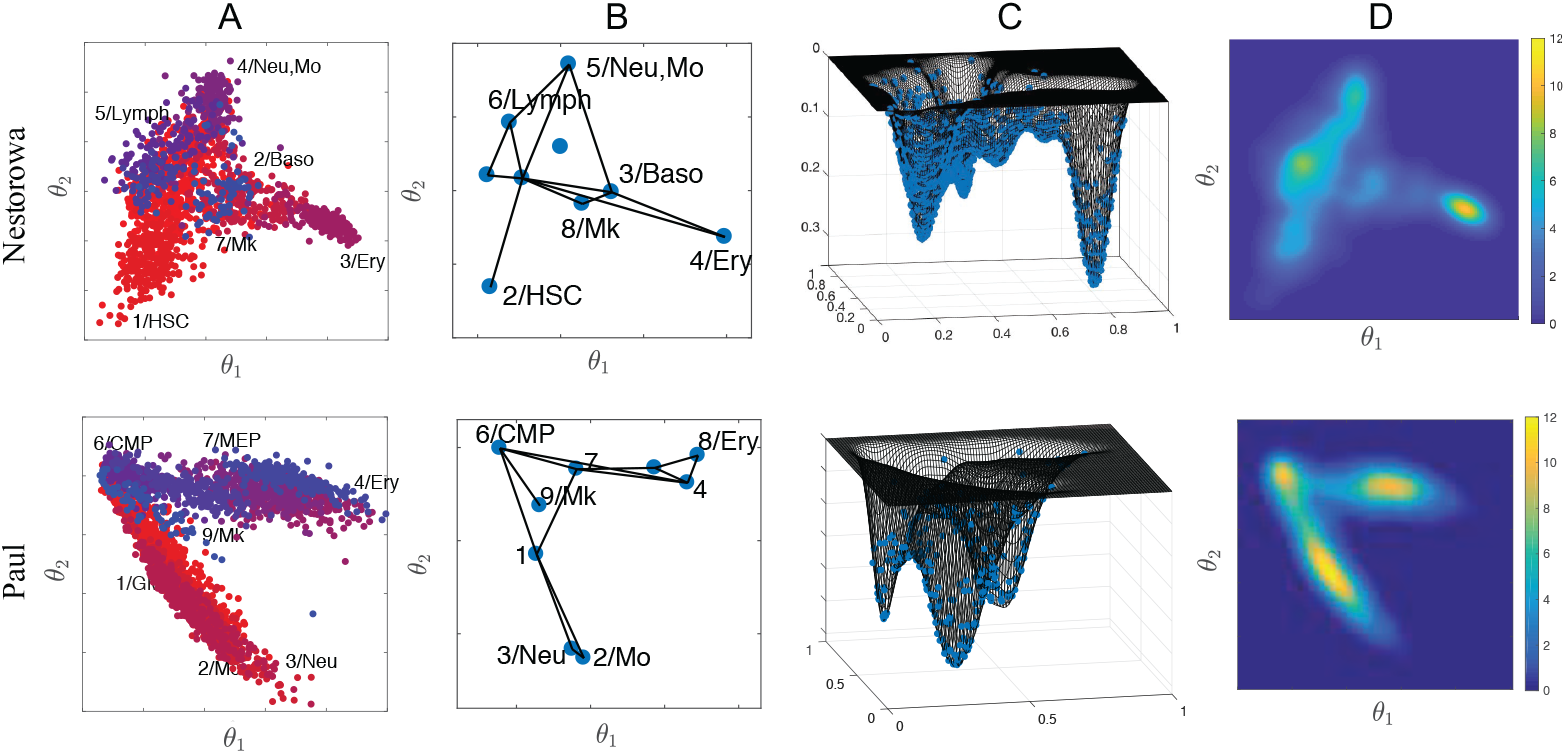
From discrete to continuum cell states. Single cell data from Nestorowa et al. (2016) and Paul et al. (2015) projected on the first two diffusion component space (A). Distinct cell types classified in the original paper are either illustrated with different colors or annotated on the graph nodes. The corresponding graph (B) and homeostatic cell distribution on the first two diffusion component space (C,D) are used to calibrate mathematical models of cell state-transitions.

The homeostatic cell distribution *u*_*s*_ on the reduced dimensional space is computed by kernel density approximation [15, 21]. The computed cell landscapes viewed in different angles are shown in Figs 2C and 2D. The cell distribution on the graph can be similarly obtained after reallocating the cells to the node, that is, the center of each cluster. The cell distribution on the continuum space provides an intuitive method to compare the relative concentration of different cell lineages, including the intermediate cell states. We observe high concentration of MEP and Ery cells that are localized at the far right (large *θ*_1_) in both data. The Nestorowa data has more diverse cells including the common lymphoid progenitors that are visible on the left (small *θ*_1_, and intermediate *θ*_2_), while the Paul data has evenly distributed cell types among the stems cells (CMP) and the two different lineages.

Let us summarize the properties of the graph and space models before we present simulation results. The cell landscape depicts the advantage and disadvantage of the two approaches. The graph model has its strength that the cell lineages between the known cell states can be more easily identified compared to the space model. The cell concentration moving toward different edges can be clearly distinguished as the cell lineages to different cell types. Accordingly, counting the number of cells in each discrete cell types is more straightforward, for instance, by integrating the cell distribution along the edges half way as in Eq (11). Although the space model has ambiguity on classifying the cells into discrete cell types, the number of cells in each discrete cell states can computed by assigning weights to integrate as in Eq (10). Moreover, we emphasize that the advantage of clear cell states in the graph model is also its limitation at the same time, since it restricts the approach to only study the presumed cell types and lineages.

The advantage of space model is its potential of exploring novel cell states that deviates from the known cell types. While the graph model cannot explore the cell states that is not already included in the graph structure, the space model can immediately study the abnormal trajectories and emergence of cells at any space location. We will show later in our simulation that the hypothesis of genetic and epigenetic alteration can be studied directly in the space model, without projecting it once more on the graph structure. Moreover, the space model is more sensitive to genetic and epigenetic variation than the graph model, although when the variation is large and a considerably distinct cell type arises, the graph model can append another cluster node.

In the following sections, we consider mainly two application problems, namely, normal hematopoiesis and abnormal hematopoiesis differentiation, resulting in myeloid leukemia as example applications of our modeling approach.

#### Calibrating the mathematical models to normal hematopoiesis

We demonstrate that normal hematopoiesis can be visualized by both models on the graph and on the space of two-dimensional diffusion component. See S1 Fig and S2 Fig for the advection and reaction terms used in the space model. Fig 3 shows a cluster of stem cells differentiating in to the entire cell states on the graph and reduced space using Nestorowa data [19] and Paul data [20]. The initial condition is imposed as approximately 10% of cell capacity in normal condition mostly composed with stem cells. On both graph and space models, nontrivial amount of most matured cell types, particularly, Ery and Neu/Mo cells arise around *t* = 30, and further recovers the full landscape after approximately *t* = 50. In particular, the observation that the matured cells quickly proliferating to fill in the capacity agrees in both simulations from Nestorowa and Paul, while the effect is more significant in Paul’s data due to shorter distances of the matured cell states from the initially administered cells.

**Fig 3.**
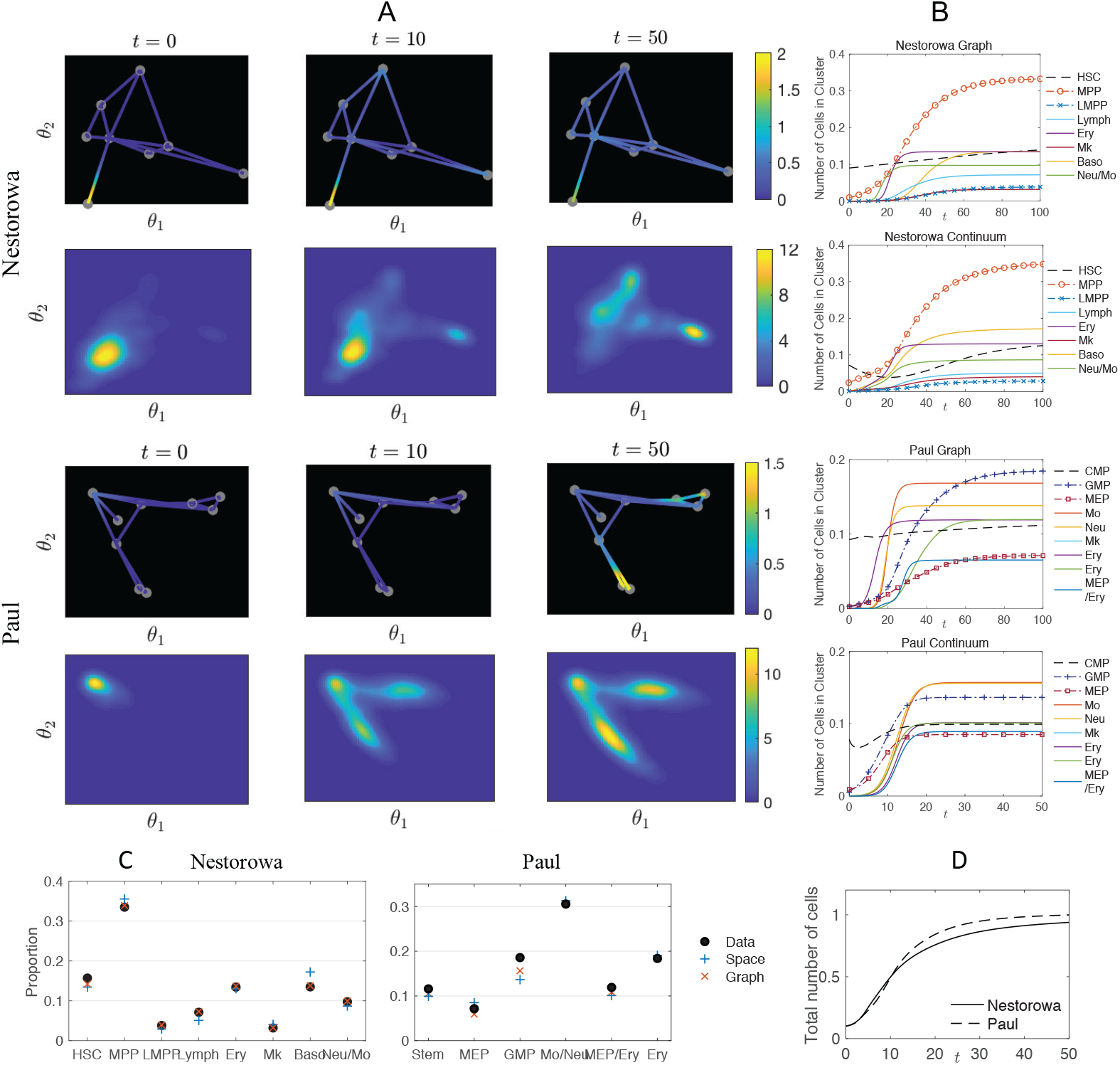
Dynamics of cell distribution during normal hematopoeisis. Evolution of cell state density *u*(*t, x*) on the graph with 8 to 9 nodes, and *u*(*t, θ*) on the diffusion component space is shown during normal hematopoeisis using single cell data from Nestorowa et al. (2016) and Paul et al. (2015) (A). The initial stem cells differentiate to progenitors and more matured cell types and recover the entire cell landscape. Numbers of cells in each type/cluster using the space model and the graph model (B) are successfully calibrated to the observed data so that at *t* = 100 each model predicts the correct cell ratios to within ±5% (C,D). We remark that the full capacity using Paul data is reached more rapidly compared to the Nestorowa data, due to its shorter distance in the reduced space.

The advantage of graph model is apparent that we can observe distinct cell states as a mass of cells shifting from a node to distinct edges toward different cell types. For instance, the cells differentiating from the MPP cluster to either Neu/Mo lineage and Ery lineage can be clearly separated in the graph models, while those can be ambiguous in the two-dimensional distribution. Still, we can compute the number of cells in each discrete cell types in both models as shown in Fig 3B. We observe that the number of cells reaches the full capacity at later times around *t* = 100, with the intended ratio of cell numbers in each discrete cell types approximating the given data [19] in Fig 3C. We comment that the recovery is more rapid for larger values of *ν* and larger number of initial stem cells *ρ*(0) (see S8 Fig).

#### Using the model framework to simulate Acute myeloid leukemia (AML) pathogenesis and progression

In this section, we once more compare the graph and space models with an application to abnormal differentiation under leukemia pathogenesis and progression. We first consider AML model in the context similar to the previous section that involves known progenitor cells that exemplifies the advantage of graph models. However, we will further show how aberrant differentiation and epigenetic plasticity of leukemia pathogenesis motivates the spatial model.

AML results from aberrant differentiation and proliferation of transformed leukemia-initiating cells and abnormal progenitor cells. We model AML pathogenesis based on known behavior of a genetic knock-in mouse model that recapitulates somatic acquisition of a chromosomal rearrangement, inv(16)(p13q22) [22, 23], commonly found in approximately 12 percent of AML cases. Inv(16) rearrangement results in expression of a leukemogenic fusion protein CBF*β*-SMMHC, which impairs differentiation of multiple hematopoietic lineages at various stages [24–26]. Most notable in such leukemia pathogenesis and progression is the increased in abnormal myeloid progenitors, which has an MEP-like immunophenotype and a CMP-like differentiation potential [25]. Experimental studies [27] show that such MEP attains a predominant increase in pre-megakaryocyte/erythroid (Pre-Meg/Ery) population (ranging from 5 to 12 fold) accompanied by impaired erythroid lineage differentiation [28]. This refined phenotypic Pre-Meg/Ery population consists partly of the CMP and MEP populations which are identified using conventional markers [19, 29].

In our model, the abnormal leukemic progenitors are regarded as intermediate cell state along the edges connecting CMP (or MPP) and MEP (and Ery) in the graph model, and the corresponding locations in the space model. We assume a 10 fold increase on average in those population by lowering *d*(*θ*) and *d*_*k*_(*x*) in Eq (4) and (7) that controls the local cell capacity. In addition to the over-proliferation, another aspect of the leukemia pathogenesis of our interest is the impaired differentiation of erythroid lineage differentiation, where it can be modeled by reducing the cell differentiation *V* (*θ*) in Eq (5) and *V*_*k*_(*x*) in Eq (1) toward Ery (See S1 Appendix and [12] for the details). The corresponding results are shown in Fig 4, where we modify the model to leukemia progression at *t* = 10. The cell distribution changes from the normal cell landscape at *t* = 10 to an increased population of Ery (MEP) and nearby cells at *t* = 20 in both graph and continuum model.

**Fig 4.**
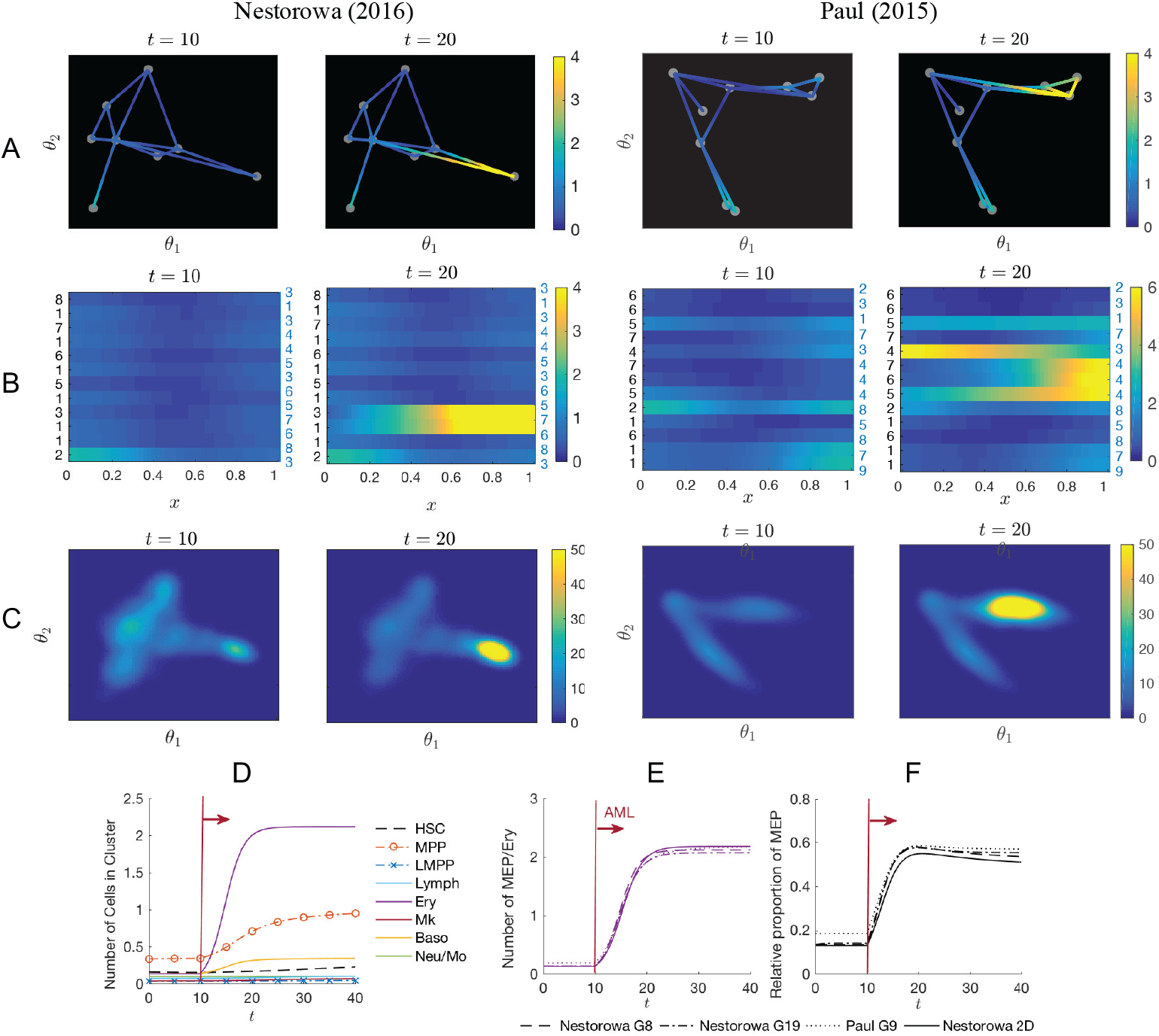
Predicting abnormal cell differentiation during leukemia progression. Cell distributions during leukemia pathogenesis and progression are shown on the graph model (A, B) and space model (C) show the effect of over-proliferation and differentiation block in the myeloid lineages. In particular, we observe the blow up of cell states near MEP and Ery in both Nestorowa and Paul data. After the initiation of AML at *t* = 10, MPP and Ery (CMP and MEP) increases using the space model and Nestorowa data (C). The number of leukemic Ery cells show a 10 fold increase within two weeks in both graph and space models (E, D), and the relative number of leukemic Ery cells comprises 50–60% of the total population (F).

We observe a 10 fold increase in the Ery (MEP) population, which includes the abnormal myeloid progenitors, in both graph and space model across the data sets as shown in Figs 4D and 4E. The blow up occurs within two week period, that shows the rapid expansion of the leukemic cells. We observe rise in the MPP (CMP) cluster as shown in the results from Nestorowa data, that is similar in Paul data as well. The total proportion of leukemic cells comprise 50–60% of the total population.

In our leukemia pathogenesis simulation, we focus on studying the leukemic cells as a variation of cells classified using conventional markers. In this case, the graph model can interpret and include the dynamics of such cells, as well as the space model. While the simulation outcome between the graph and space model is similar, the graph model is computationally more efficient due to the fewer number of unknowns as compared to the two-dimensional space model. However, to study the abnormal cell states that may appear far away from the known or existing landscape, the space model has more freedom to include those new cell states and disrupted trajectories. In particular, abnormal regulation of cells can arise through mechanisms outside of gene transcription such as epigenetic alterations or DNA mutations. In the following section, we study the impact of perturbation of genes in the graph and space model, particularly focusing on alterations of genes known to be involved in leukemia pathogenesis.

### Mathematically perturbing gene expression

In this section we investigate the sensitivity of altering specific genes in a prescribed manner and the impact of this perturbation in the graph and reduced space models. We keep our focus on leukemia pathogenesis and progression and consider alterations of 38 genes that are reported to be related to leukemia stem cells [30, 31], although we emphasize that these genes serve simply as examples, and are not intended to model the precise biological process of AML pathogenesis. The *j*-th gene expression level of *i* cell, 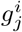, is modified as 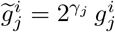, where *γ*_*j*_ is the log_2_-fold change compared to the normal cells, that ranges from −3.5 to 2.7. The full list of altered genes and magnitudes *γ*_*j*_ are shown in S1 Table. In addition, we consider the extreme case of gene alteration as the maximum level 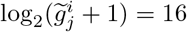 for up-regulated genes and 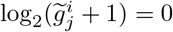 for down-regulated genes. Fig 5 shows examples of the gene expression levels that we modify in log scale including the up-regulated genes, GPR56, GATA2, and MZB1, and the down-regulated genes, LGALS3, LY86, and ANXA5. The given single-cell data in normal condition is plotted together with the altered levels of cells in regular leukemia pathogenesis and extreme level of progression. Although the extreme alteration is unrealistic, we consider such case to illustrate an example where the graph abstraction and dimension reduction algorithm clearly distinguishes the leukemic cells. In the following section, we compute the graph and reduced space using the mixture of the leukemic and normal single-cell data, and compare it with those in normal condition.

**Fig 5.**
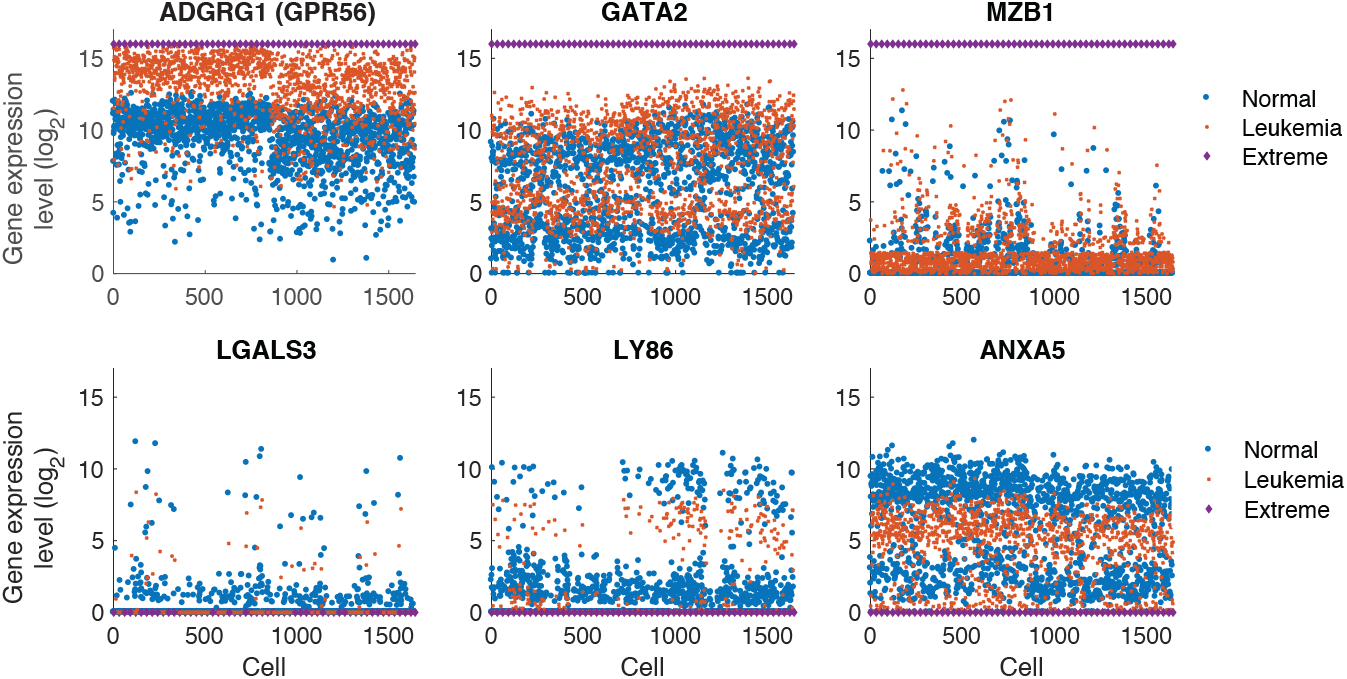
Perturbing genes associated with leukemia stem cells. Examples of expression levels of genes (in log_2_ scale) that are associated with leukemia stem cells and pathogenesis, including up-regulated GPR56, GATA2, and MZB1, and down-regulated LGALS3, LY86, and ANXA5. We show these subset of genes simply to illustrate the process. The normal single-cell data (blue circle) and modified gene expression (red square) are shown together, with the case of extreme levels (purple diamond).

#### Effect of perturbing genes on graph and reduced component space

We first study the sensitivity of the reduced component space using diffusion mapping [13]. Fig 6 compares the altered leukemic single cell data in the normal reduced space (*θ*_1_, *θ*_2_) and the recomputed reduced space using both normal and leukemic cells, denoted as 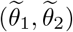. We also denote 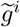 as the altered gene expression level of the normal single-cell data *g*^*i*^. The left-most column shows the projected leukemic single cell data 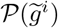 in the normal reduced space, where the leukemic cells are located toward the left-bottom compared to the normal cells in Nestorowa data, and upwards in Paul data. The effect of gene modification is shown more clearly in the presented vector field 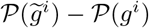. The recomputed reduced component space with both normal and leukemic data show similar directional trend, but with slightly larger magnitudes. The reduced space computed with the altered cell data to extreme levels are shown in the right-most column, and they distinctly separate the two cell clusters indicated the second diffusion component *θ*_2_.

**Fig 6.**
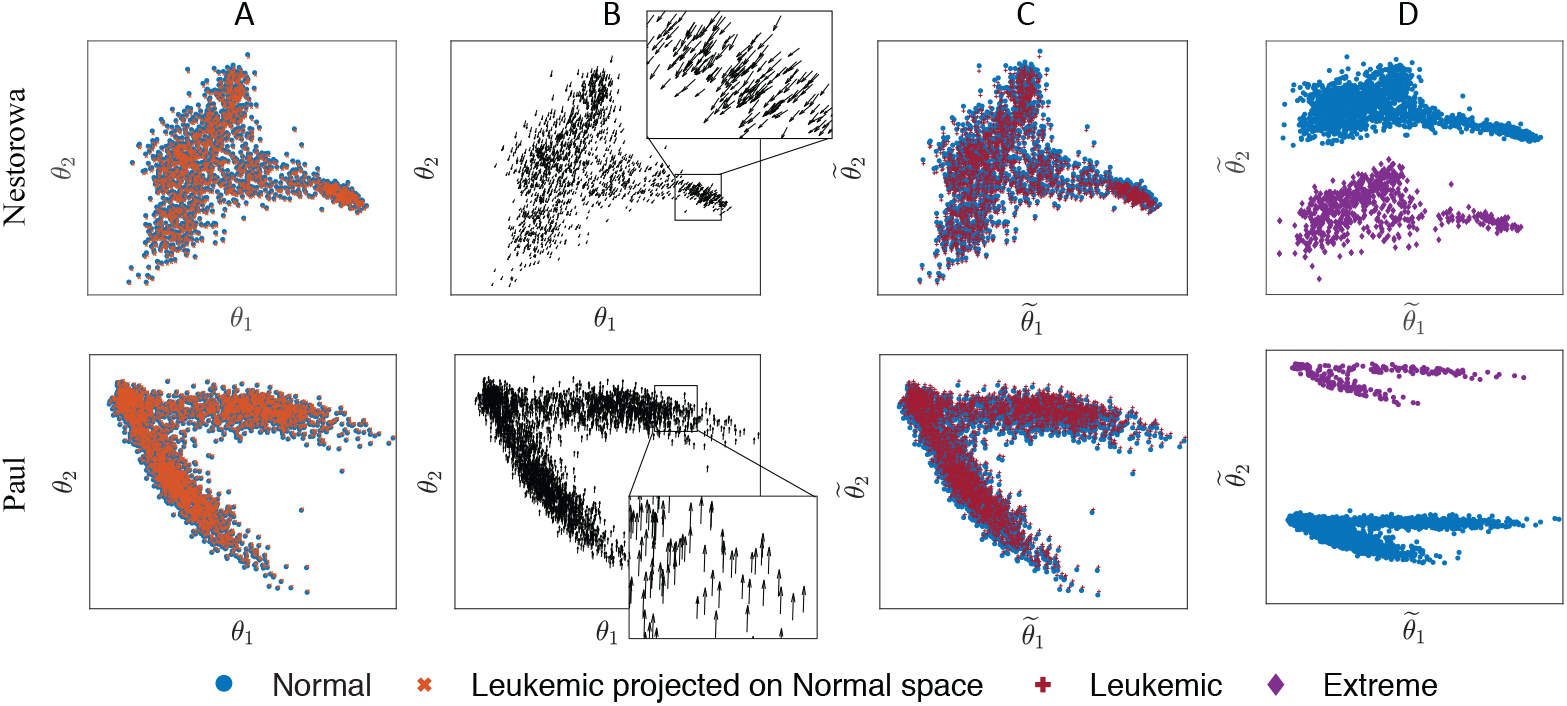
Effects of perturbing genes on cell state-space. Projection of perturbed leukemic single cells on the normal diffusion component space (A), and their altered directions (B). The reduced diffusion space with both normal and leukemic cells (C), and normal and extreme cells (D) show similar relationship between two cell clusters, however with larger separation. The top figures are computed with Nestorowa data [19] and the bottom figures with Paul data [20].

Similarly, we study the impact of leukemic gene perturbation in graph abstraction using partition-based graph abstraction [17]. Fig 7 shows the clustered cell types and the corresponding graph using both normal and leukemic cell data, and with data in extreme condition. The presented results are with Nestorowa data. The clustered cell types and leukemic cells are annotated to depict the cluster properties, and in case of regular leukemia progression with cells altered in regular magnitude, we observe that there is no cluster that separates the leukemic cells. Thus, the information of leukemic gene alteration is lost within the clustering algorithm, and the model on such abstracted graph is not capable of studying the perturbed cells. The cells are clustered into separate nodes in the extreme case of modifying the gene levels to their maximum and minimum, however we comment that this is an unlikely scenario. See S4 Fig for other magnitudes of perturbation and Paul data.

**Fig 7.**
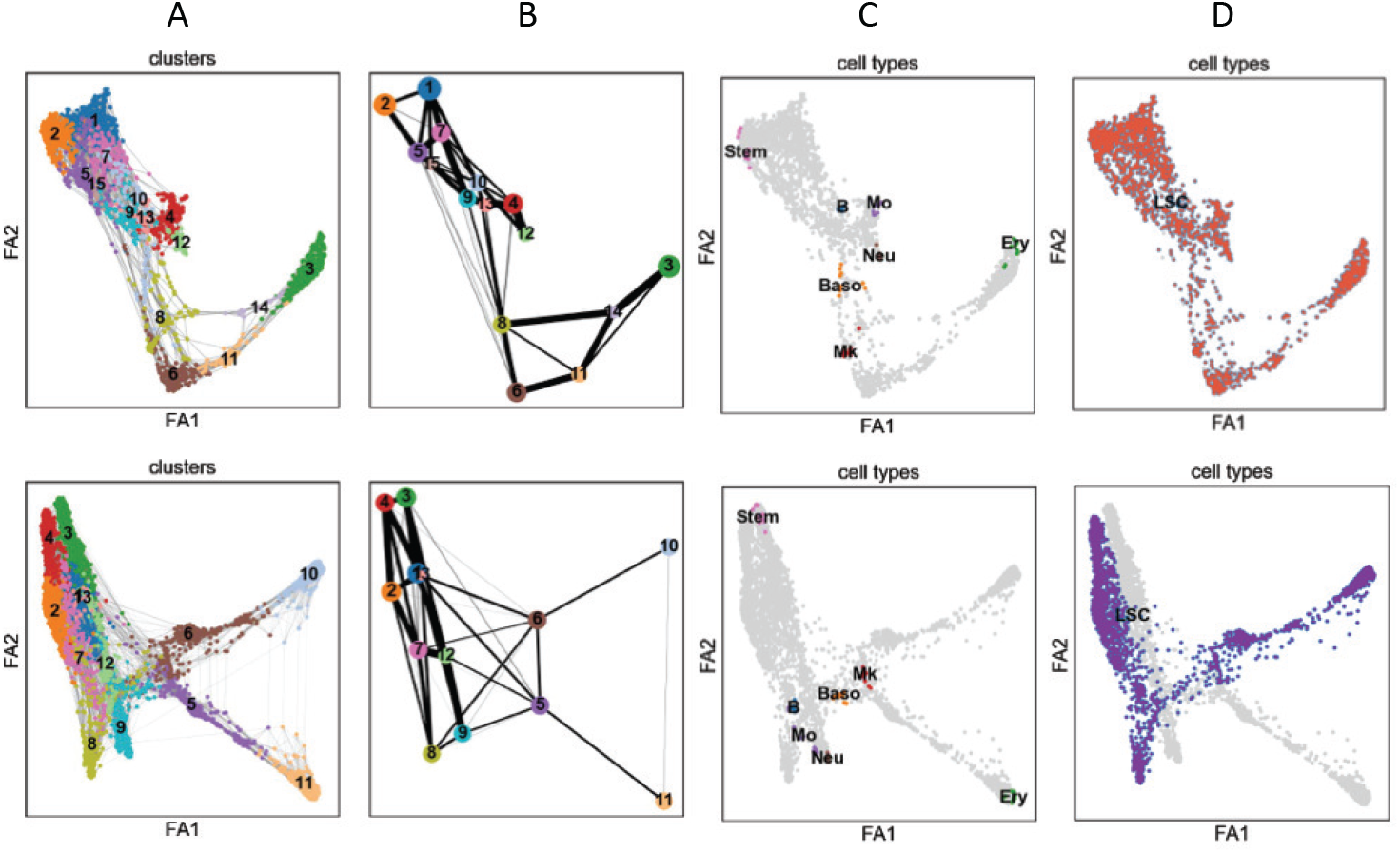
Effects of perturbing genes on abstracted graph. Graph computed from normal and perturbed leukemic cells (top), and from normal and extreme condition (bottom). The shown figures are (A) cell clusters, (B) abstracted graph, (C) annotated normal cell types, and (D) leukemic cells. The graph algorithm does not distinguish the perturbed leukemic cells in regular magnitude to the normal cells, so that the perturbed information is lost. The algorithm distinguishes the leukemic cells when the data is modified to the extreme values of gene expression level although the data is unrealistic.

A strength of the continuum cell state-space model is its capability of interpreting perturbation of gene expression levels or new incoming cell data regardless of its relation to the primary data (Figs 6 and 7). As shown in our results, the leukemic alteration is successfully projected in the reduced space, while the abstracted graph lost the information. Although the projected directions in the reduced space can be once more projected on the graph, it does not make sense to do so when the direction is orthogonal to the edges. The space model has its advantage especially in this case, where the projected direction of cell states can be directly implemented.

The remaining question is how to interpret new cell states in the space model that may arise far away from the cell states that can be identified by conventional markers. Hence, we propose some measures in Eqs (12)–(13) to guide the interpretation. Fig 8 shows an example of the rescaled correlation quantities *r*_*f,j*_ and *r*_***v**,j*_ computed with Nestorowa data. The first row show results of the correlation *r*_***v**,j*_ to the average leukemic directional vector ***v*** = (*−*0.068, − 0.206). The gene expression levels of genes that have large values of *r*_***v**,j*_ are depicted in the figure, namely, PLAC8 and CAR2. We remark that those genes have strong local correlation 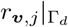 on Γ_*d*_ = {0.3 ≥ *θ*_1_ ≥ 0.9, 0.3 ≥ *θ*_2_ ≥ 0.5} as well. Fig 8 shows the correlation of all 3991 genes, where the red bars highlight the leukemia related genes we modify (S1 Table) and we observe large magnitudes in some of the genes. The second row shows the correlation *r*_*f,j*_ to a specific cell state at the reduced space, where we choose *θ*^∗^ = (0.5, 0.35), which is approximately an intermediate location between MEP and CMP cells, and 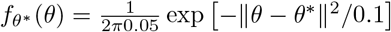. APOE and CLEC12a genes show the largest magnitude of *r*_*f,j*_, and similarly, we can identify the leukemia related genes that show strong correlation to cell state *θ*^∗^. Although more careful and rigorous approach should be developed to characterize the new arising cell states, *r*_*f,j*_ and *r*_***v**,j*_ provides an efficient method of initial screening of possible related genes.

**Fig 8.**
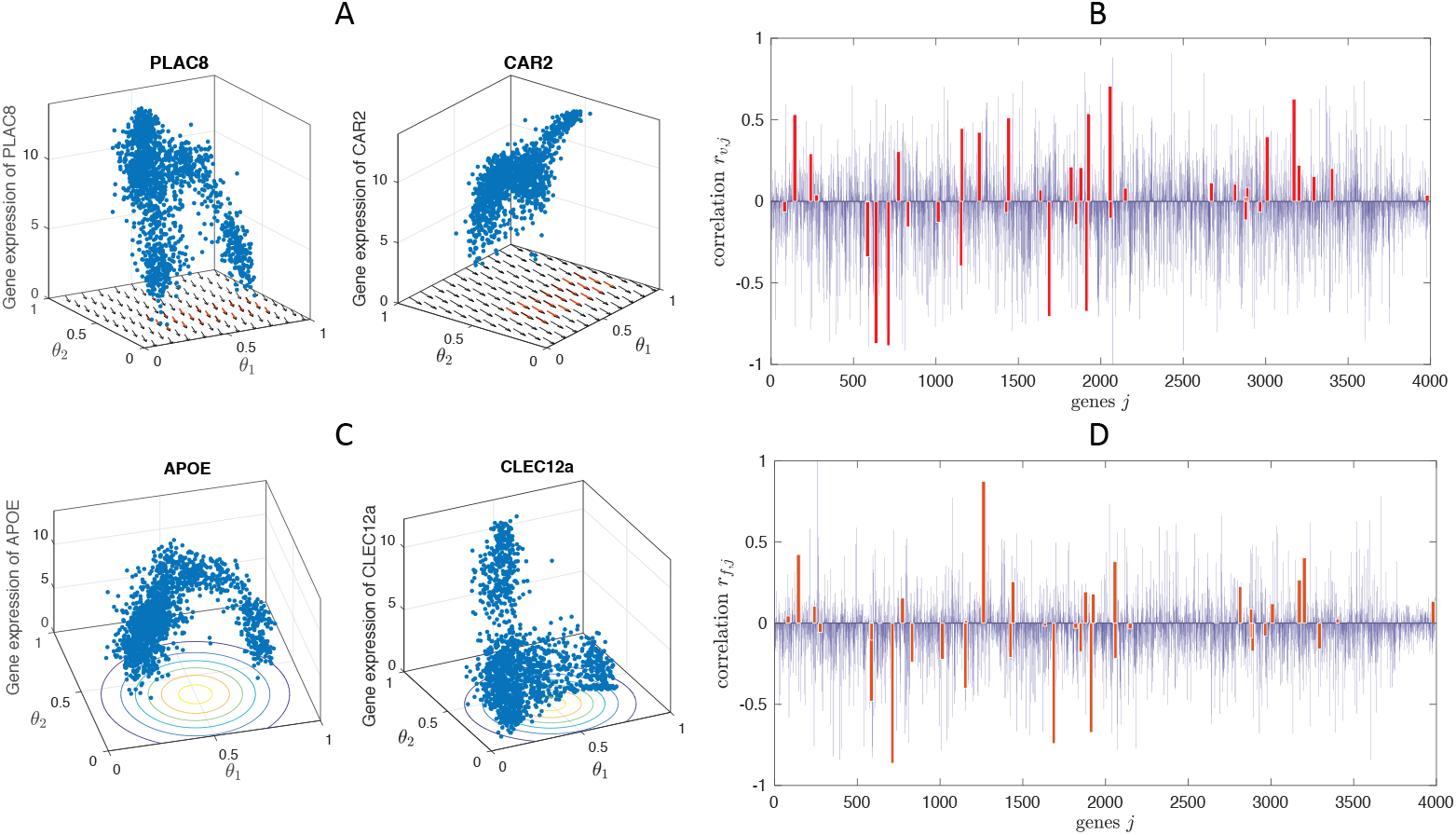
Interpretation and mapping of model-predicted novel cell states. In order to identify novel cell types predicted by the mathematical model, gene expression levels 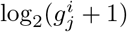 that are strongly correlated to the direction of leukemic alteration ***v*** = (*−*0.068, −0.206) (A), and to the reduced space location *θ*^∗^ = (0.5, 0.35) (C). The rescaled correlation *r*_*f,j*_ (B) and *r*_**v**,*j*_ (D) computed for all the genes in Nestorowa data are shown, and the leukemia related genes are marked in red bars.

#### Simulating AML pathogenesis by perturbing known leukemia genes

In this section, we incorporate the perturbed leukemic gene data in the AML simulation using the space model. In particular, we are interested in studying the impact of leukemic genes alteration on the cell distribution during AML progression. We compute the abnormal differentiation of leukemic cells by projecting the altered single cell data of MEP and Ery cells to the normal diffusion component space as it is shown in Fig 9. The aberrant differentiation vector 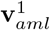 shows the shifts of cells toward the location where no cell data occupies. Therefore, in addition to modifying the advection term according to the altered gene data, we assume an emergence of new abnormal cell state. In particular, we take the cell state location at *θ*^∗^ = (0.610, 0.215) in Nestorowa data, and at *θ*^∗^ = (0.6, 1.0) in Paul data, and use Gaussian functions centered at *θ*^∗^ to obtain 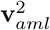. The corresponding vector fields are also shown in Fig 9.

**Fig 9.**
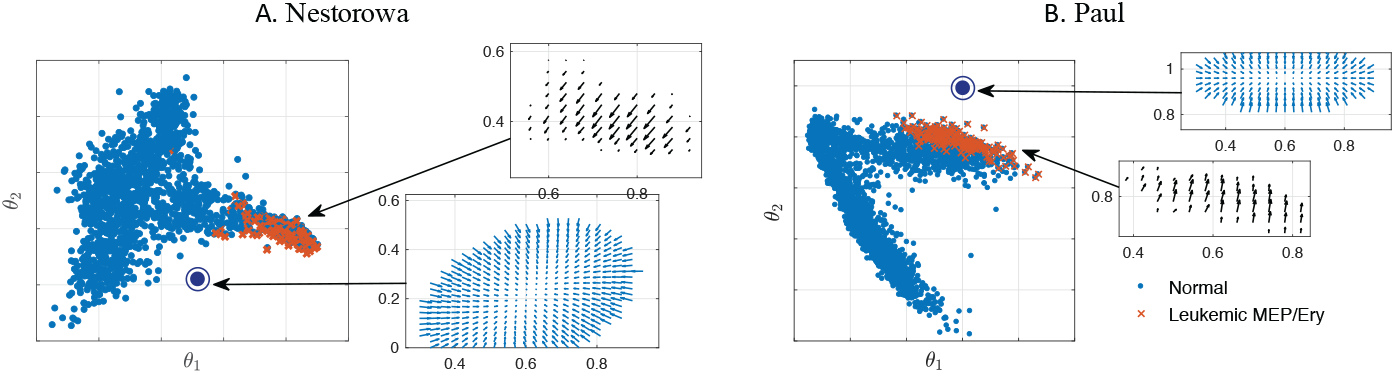
Modeling AML pathogenesis and progression by perturbing cell states directly in the cell state-space. The direction of abnormal cell differentiation 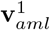 is computed from the projection of altered leukemic MEP and Ery cells (*×*) to the normal diffusion component space. Alternatively, we assume a source of abnormal cell state (⊙) at *θ*^∗^ = (0.610,0.215) in Nestorowa data (A) and at *θ*^∗^ = (0.6,1) in Paul data (B) to model 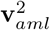.

For AML progression, the advection term is modeled with the prescribed vector field as 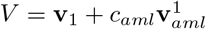 or 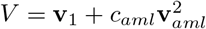, where *c*_*aml*_ parameterizes the perturbation magnitude. We further perturb the model by increasing the proliferation of the new leukemic cells at *θ*^∗^ by appending 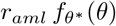 to *R*, where *r*_*aml*_ parameterizes the over-proliferation. The cell distribution *u*(*t, θ*) with 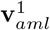 and 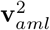 for different values of *c*_*aml*_ = 1, 2, 10 are presented in Fig 10 and S6 Fig. In the cell landscape with 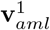, we observe increased MEP cells and abnormal progenitors arising in the direction of left-bottom, especially for large values of *c*_*aml*_. The model with 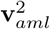, a new cell type further down in the cell space emerge and dominates the population. With the model 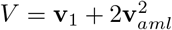, new abnormal cells appear around *t* = 10 and dominates the population at *t* = 30. The total number of cells and cell number in each discrete cell type is plotted in Fig 10, where the effect of large values of *c*_*aml*_ and *r*_*aml*_ is more clearly shown. The total number of cells increases more than 10 times the initial size after *t* = 30 when *c*_*aml*_ = 10 and *r*_*aml*_ = 0. When the over-proliferation term is appended as *r*_*aml*_ = 1, the total number of cells increases more rapidly, for example, up to 100 times the initial size and the number of cells in most of the myeloid lineage increases.

**Fig 10.**
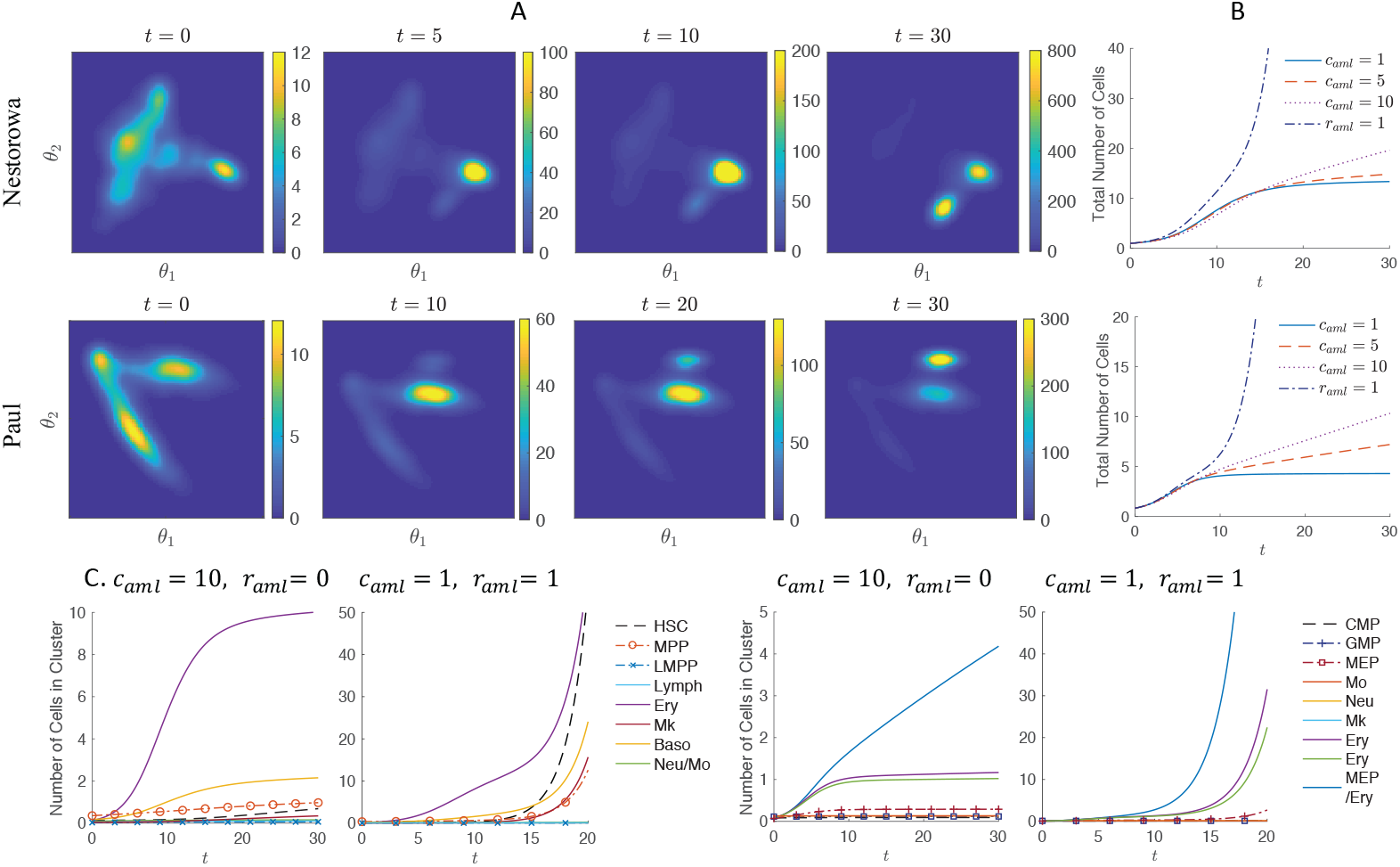
Cell state-transition dynamics during leukemia pathogenesis and progression. The evolution cell state distribution *u*(*t, θ*) with *c*_*aml*_ = 2 (A) and total number of cells (B) in AML is shown using model 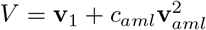. More rapid progression of AML in terms of cell number is observed for larger values of *c*_*aml*_ and *r*_*aml*_. The number of cells in each cluster (C) shows the increasing level of leukemic Ery cells, and the effect of perturbing the proliferation *r*_*aml*_ > 0 that it facilitates the AML progression.

### Code availability

The simulation codes are available from github.com.

## Discussion

We have shown how to construct and calibrate mathematical models of cell state-transitions using single-cell sequencing data. We compare two approaches: solving equations on graphs and solving equations on a continuous cell state-space. Each approach has strengths and limitations. Selecting an approach for a given application or dataset will depend on the type of biological data and the nature of the scientific question.

When the modeling application and quantity of interest includes well-known cell lineages and relation between discrete cell states, the graph model is more appropriate due to its ability of distinguishing distinct cell lineages more clearly compared to the continuum space model. Dynamics of cell number in discrete cell types, alteration of proliferation and apoptosis in particular cell type, differentiation block, and emergence of intermediate cell states can be straightforwardly quantified and studied. However, to explore the cell states beyond the known cell lineages, the continuum space model is more advantageous since it includes all pathological and intermediate cell states, rather than confining the model into presumed cell lineages. Moreover, the continuum model can incorporate a relatively small genetic and epigenetic alteration that the graph abstraction may not recognize, and study abnormal trajectories that yield unconventional cell states.

We selected genes to perturb to simulate AML based on genes known to be associated with leukemia pathogenesis. We do not intend for this to be an accurate model of the biological process, rather, as an illustration of how one may select sets of genes and perturb them in a prescribed fashion in order to study the effect on cell state-transition dynamics. This approach assumes that AML pathogenesis originates from changes in gene expression in specific cell subsets, which is limited by our identification of these genes based on literature. We acknowledge this is a limitation of the modeling approach, although we also note that our model predictions are consistent with known features of AML progression.

### Comparison to other approaches

Although at the time of this work there are relatively few mathematical models published which utilize single-cell sequencing data, there are a few notable exceptions. Of particular note are works which use modeling and simulation to generate synthetic *in silico* gene expression datasets [32]. These important approaches to mechanism based mathematical modeling may also be used to study and predict the effects of perturbations on cell state distributions. They may also be used as computational controls to benchmark analysis tools and potentially to benchmark and compare mathematical models, although using a model to benchmark other models can lead to consistent but incorrect circular reasoning and caution is warranted. Another example is Ferrall-Fairbanks and Papalexi et al, who use mathematical analysis to generate novel quantifications of cell heterogeneity in cancer or immune cell subsets respectively [33, 34]. These methods may be used to map and interpret novel cell states predicted by mathematical models or similarly as a method to interpret model-predicted changes in cell heterogeneity following a perturbation.

Schiebinger et al compute and predict differentiation trajectories in cell development using optimal transport (OT) [35]. This approach considers the optimal transport of cells as a mass flowing along differentiation trajectories, and is conceptually the most similar to our approach. As presented, Schiebinger et al do not use the OT framework to examine perturbations of cell states or genes along the differentiation trajectory, although this is possible with an OT model. Setty et al present a method to compute cell fate probabilities [36], which may also be achieved by inferring cell state-transition dynamics with lineage trees [37]. Fischer et al have demonstrated a method for inferring population dynamics from single-cell sequencing data [38], Sharma et al use longitudinal sequencing to study drug-induced infidelity in the stem cell hierarchy [39], and Karaayvaz et al show how to use single-cell sequencing to examine drug resistance in breast cancer [40]. These approaches and analysis methods may be used to inform and potentially calibrate mathematical models of cell population dynamics or response to treatment-induced perturbations.

Recently, vector fields derived from RNA velocity [41] have been used to infer potential energy or fitness landscapes for cell state-transitions [42]. These approaches may be used to inform the computational domain for mathematical models that we present here, however, an important limitation of the RNA-velocity approach is extrapolation of the vector field outside of the data range. This underscores the need for hypothesis-based and model-guided approaches to inform the shape of these fields. This limitation also applies to the rapidly growing field of deep learning approaches [43] to analyze single-cell sequencing data, namely, whether the learning algorithm can effectively make predictions to datasets which are not sufficiently similar to those upon which it has been trained. We believe that the future likely involves a merger of mathematical modeling with machine learning, in which mathematical models are used to inform learning approaches and impute sparse data as has been recently shown by Gaw and Rockne et al [44, 45].

## Opportunities and limitations of modeling with single-cell sequencing data

There are pros and cons, opportunities and limitations to mathematical modeling with single-cell sequencing data. The advantages and potential opportunities include: a wealth of available data, richness and complexity of each data set, a focus on the cell level, opportunity to study dynamics in hierarchically structured state-based relationships between cells, and an ability to perturb individual cells and/or genes within cells to predict dynamics of state-change at cellular level. The most significant strength of mathematical modeling is the ability to use and generate hypotheses that may not be directly evident from the data; for example, extrapolation of RNA velocity fields beyond the dataset boundaries or to interpret and predict novel cell types which may not otherwise be clearly identified with known canonical cell state markers. Another advantage of our approach is the use of pseudo-time analysis of data collected at a single timepoint to calibrate the models, however, the models can also be calibrated directly to time-sequential single-cell datasets, which we expect to become more commonly available as single-cell sequencing continues to be used as a tool to study cell dynamics.

The disadvantages and limitations include: the potential for misleading or incorrect inference due to poor data quality including drop-outs, small non-representative samples of large heterogeneous populations, no physical or micro-environmental context, no direct or physical interactions between cells, and the possibility of model predictions to be sensitive to methods of dimension reduction, graph abstraction, state-space construction, and potentially sequencing platform. Sensitivity of the modeling to experimental and computational methods may be directly studied and mitigated as we have shown in this work, however this remains potentially a significant source of uncertainty and variability in the modeling calibration and predictions.

In terms of computational cost, the graph model is more efficient since it is a multiple of one-dimensional cost, while the cost of implementing the space model increases exponentially as the dimension of reduced space increases. In our simulation, the continuum model runs approximately 10 times longer than the graph model with 8 nodes, therefore, the space approach will be reasonable when the reduced component can be truncated at two- to three-dimension, unless the numerical method is carefully implemented.

### Future work and applications

Future applications of this approach is to explore hypothesis in the resolution of single-cell genomics and study altered and novel cell states with genetic and epigenetic alterations in various biological systems and pathogenesis [46]. Incorporating effects of external perturbations such as therapy is our interest as well. Moreover, comparing the model prediction to sampling/sequencing of perturbed biological system is our most anticipated future work, for instance, we look forward to include scRNA-seq data of leukemic progenitor cells.

There are opportunities for further enhancements in our model in improving the model of cell landscape dynamics to accurately estimate cell transition pathways in the reduced component space, for instance, minimum action paths [6] and bifurcation [7, 47]. The model can be improved by obtaining parameter functions or mappings of biological quantities directly from single cell sequencing data, for example, more precisely infer the proliferation rate function. Also, developing methodologies to obtain reduced component space that captures desired characteristic of cell states [48] will help us explore our approach for other biological settings where cell states are less clearly characterized. Moreover, we propose to develop quantities, such as index of critical state transitions [47, 49], in the phenotype space that could be used to predict forthcoming major alterations in development and diseases. We also expect to be able to infer the potential landscape directly from the RNA velocity vector field [41, 42].

## Summary

In summary, despite the explosion of computational tools to analyze single-cell sequencing data, there have been relatively few mathematical models developed which utilize this data. Here we begin to explore the possibilities—and limitations—of dynamical modeling with single-cell data. We hope this work paves the way for mathematical models to be developed to guide the interpretation of these complicated datasets as they begin to be collected after biological perturbations (ex. cancer, treatment, altered developmental processes), sequentially over time, or sampled spatially within biological tissues.

## Supporting information

**S1 Table.** Summary of the required parameters. The following table summarizes the parameters in our model terms, *V*, *R*, *D*, *V*_*k*_, *R*_*k*_, and *D*_*k*_, and their biological meaning with the range.

| Parameters | Biological meaning | Range |  |
| --- | --- | --- | --- |
| $r(\theta)$, $r_k(x)$ | proliferation rate | [0.00215, 1] | [50–53] |
| $a(\theta)$, $a_k(x)$ | self-renewal rate | [0.1, 0.8] | [16, 54] |
| $\mathbf{c}(\theta)$ | differentiation vector | [0, 1] | |
| $\nu$ | phenotypic fluctuation | [0, 0.0027] | |
| $\bar{d}$ | apoptosis rate | 0.6925 | [54] |

**S2 Table.** Parameter values of proliferation and self-renewal rate. The following values are taken for each single-cell, 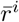 and 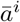, based on their clustered cell types, and then used for computing *r*(*θ*), *r*_*k*_(*x*), *a*(*θ*), and *a*_*k*_(*x*).

| | hematopoietic stem cells (HSC) ↔ progenitor cells (HPC) | | | |
| --- | --- | --- | --- | --- |
| cell type | HSC | MPP, LMPP, CMP | MEP, GMP | Neu/Mo, Ery |
| proliferation | 0.01125<br>(8.8 weeks) | 0.05658<br>(12.25 days) | 0.1612<br>(4.3 days) | 0.6931<br>(1 day) |
| self-renewal | 0.77 | 0.7689 | 0.7359 | 0.66 |

## S1 Appendix. Details of the model equation and parameters

The model terms require interpolation of single-cell data to the continuum cell state space. We use clustering to identify cell types and their cell type properties to assign those to each single-cell. S2 Table are the values we take for cluster properties. By denoting 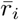 as the assigned proliferation rate of the *i*-th cluster, we compute the intermediate level of proliferation in the graph model by linear interpolation as

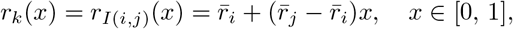

assuming that the overall proliferation of intermediate cell states change gradually. In the multi-dimensional model, we compute the interpolation based on local means as

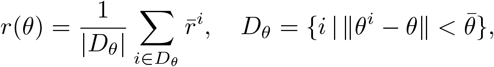

where we take 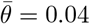. The self-renewal rate functions *a*_*k*_(*x*) and *a*(*θ*) are computed similarly. See S1 Fig for *r*(*θ*) and *a*(*θ*) computed for Nestorowa data.

To compute the multi-dimensional function on the continuum space from the single-cell data, we employ the *kernel density method* [15, 21], that is a non-parametric way to estimate the density function based on a finite data sample. Using the single-cell samples in the reduced component space, 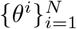, the method approximates the density function as

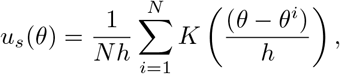

where *K* is the kernel smoothing function that we take it as a Gaussian function and *h* is the bandwidth. The optimal bandwidth to estimate normal densities can be computed by 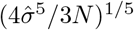, where 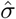 is the standard deviation and *N* is the sample size, and the optimal bandwidth for our data is computed as 0.0383 to 0.0456, however in our simulation, we choose a slightly smaller value, *h* = 0.03, to reveal more features of multiple modes. Fig 2(a,b) shows the results computing *u*_*s*_(*θ*) using Nestorowa and Paul data, and S1 Fig(a) shows the corresponding **v**_1_(*θ*).

In the diffusion term, we explore the parameter *ν* so that the phenotypic instability does not dominate the cell maturation. We compute the parameter in the range of *ν* ≤ (*L*/*T*_*d*_)^2^/4, where the distance in the diffusion space is *L* = 1 and the time that HSC differentiates to the progenitors is *T*_*d*_ = 5 ~ 30 (day), that is, *ν* ≤ 0.0027 ~ 0.01, and we consider *ν* = 0.001. Quantifying the local phenotypic instability in the reduced component space, and justifying this term is our future work.

To compute the reduced component space using dimension reduction approaches, we employ *diffusion mapping*. See [13, 55] for the detail of the algorithm. We take the cosine distance, *k*(*x*^*i*^, *x*^*j*^) = 1 − *corr*(*x*^*i*^, *x*^*j*^) for the Nestorowa data and the gaussian distance 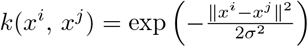 for Paul data with *σ* = 50. From *L*(*i, j*) = *k*(*x*^*i*^, *x*^*j*^), the diffusion mapping use parameter *α* to tune the influence of density of the data points as

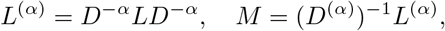

where *D*^(*α*)^(*i, i*) = Σ_*j*_*L*^(*α*)^(*i, j*), and we choose *α* = 0.5. From the eigen-decomposition of *M φ* = *λφ* and ordered eigenvalues 1 = *λ*_0_ ≤ *λ*_1_ ≤ *λ*_2_ ≤ …, the corresponding right eigenvectors, *φ*_1_, *φ*_2_, … are the diffusion components. We truncate the reduced space at the second diffusion component, where the eigenvalues are *λ*_1_ = 0.1039, *λ*_2_ = 0.0326, *λ*_3_ = 0.0167, *λ*_4_ = 0.0135 for Nestorowa data, and *λ*_1_ = 2.4653e-03, *λ*_2_ = 5.8338e-04, *λ*_3_ = 9.7792e-05, *λ*_4_ = 7.0364e-05 for Paul data. For a comparison of diffusion mapping to two-dimensional reduced component space using other dimension reduction algorithms, see S3 Fig.

For the pseudotime inference, we use the algorithm developed in [56]. The diffusion distance between two cells are computed as

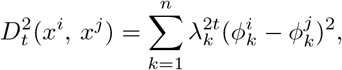

and the pseudotime distances are computed based on this distance. We choose three extreme points in each of the three clusters, stem cell, Ery, and Neu cell types, that are the furtherest in the diffusion component space, and infer the lineage between the extreme cells. After computing the pesudotime of each single-cell we compute the local average direction to the neighborhood cells that are in later pesudotime similar as in the computation of local means for *r*(*θ*). The computed results are shown in S2 Fig(b) with the interpolated vector at the grid points, S2 Fig(c).

**S1 Fig.**
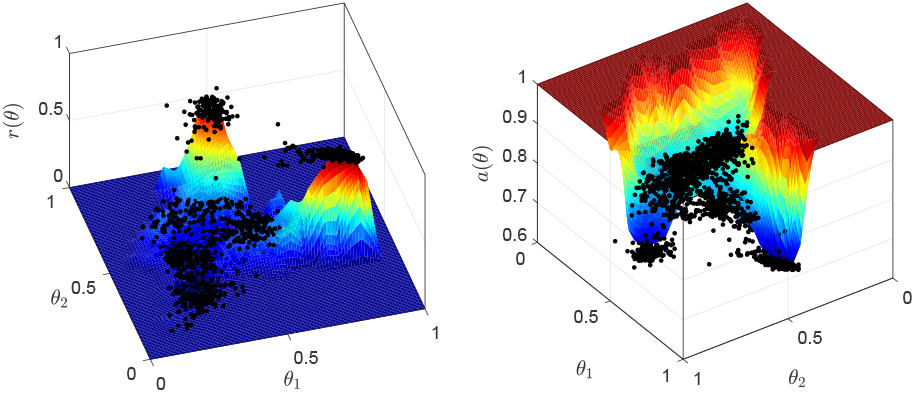
Cell proliferation rate *r*(*θ*) and self-renewal rate *a*(*θ*) computed from the single-cell data. The black dots are the rates of data.

**S2 Fig.**
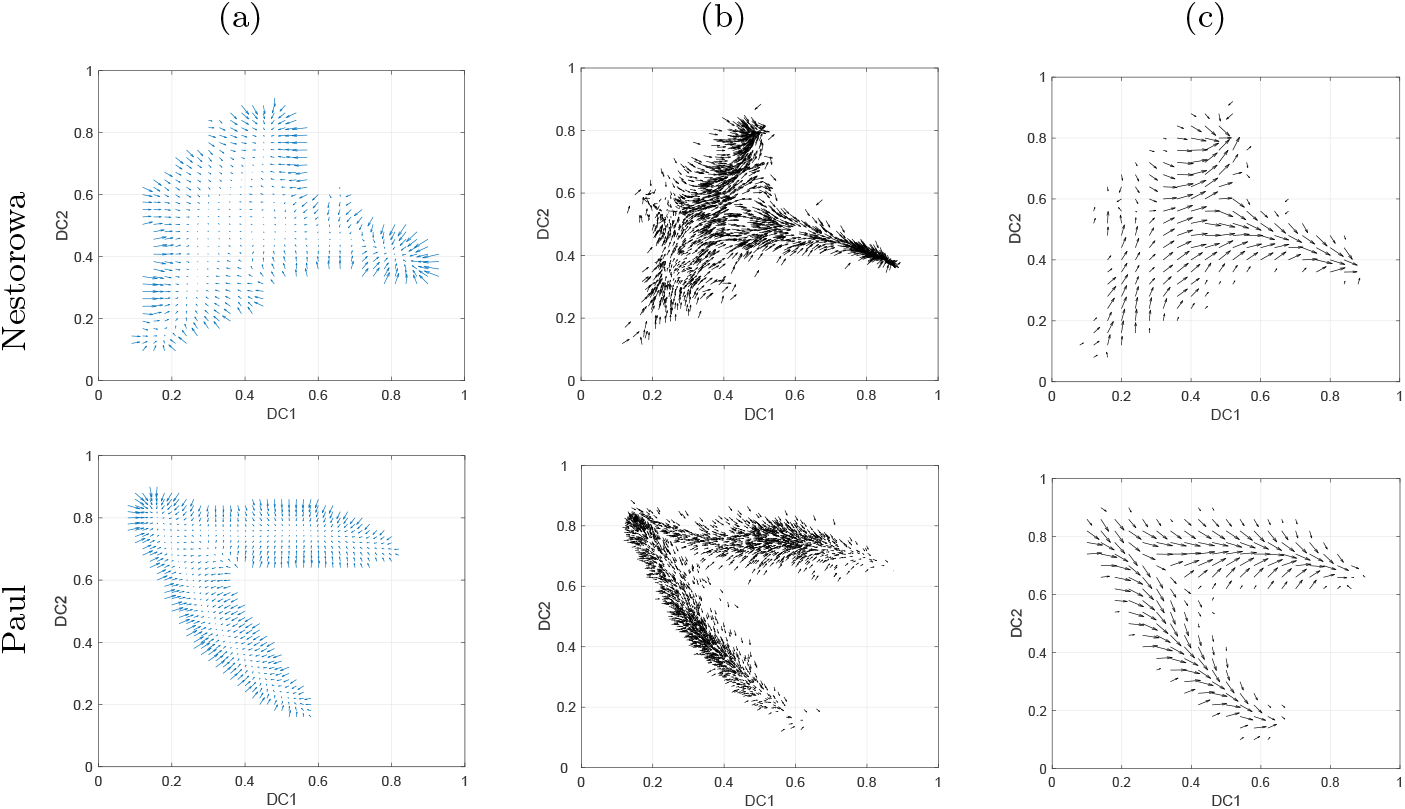
Pseudotime dynamics. The homeostasis cell differentiation vector **v**_1_ (a), and the direction of active cell differentiation obtained from diffusion pseudotime analysis (b), and the pseudotime vector interpolated at the grid points **v**_2_ (c) are presented. We remark that, **v**_2_ corresponds to the cell differentiation along the edges in the graph model.

**S3 Fig.**
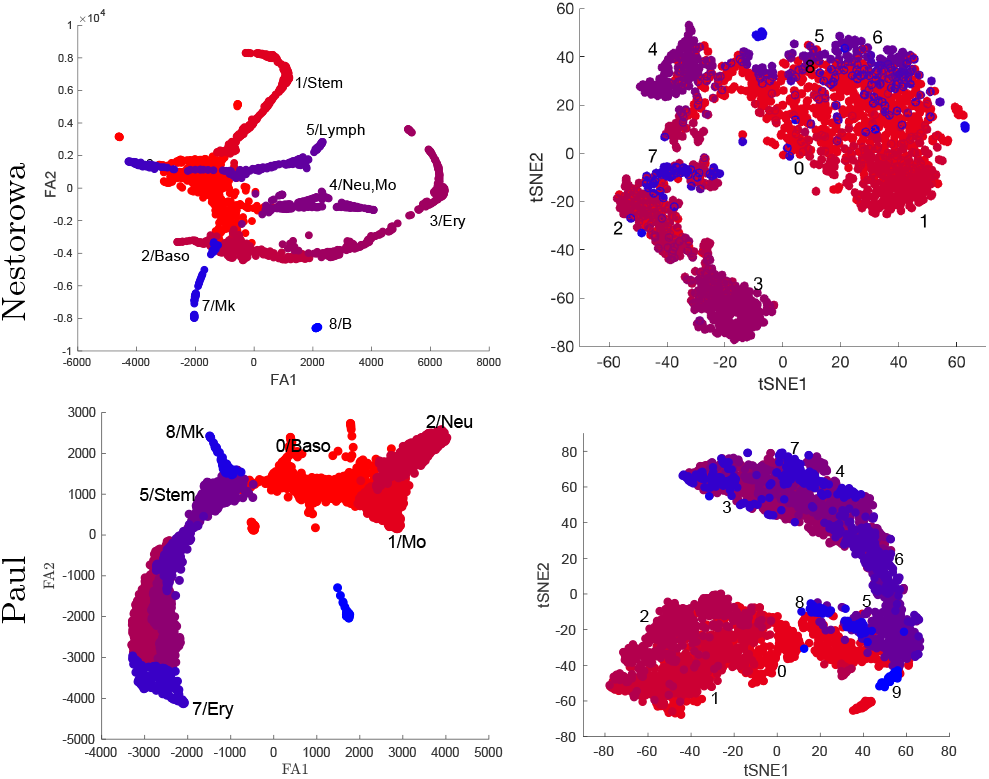
Comparison of dimension reduction methods. Dimension reduction algorithms that focus on preserving local structure are not appropriate to infer global trajectory. Compare the following results of single cell data projected on the reduced component space of ForceAtlas2 (FA) [57] and t-stochastic neighbor embedding (tSNE) to diffusion component space in Fig 2.

**S3 Table.** Gene alterations in Leukemic stem cells. From the genes that are reported in [30, 31], we find all the gene that are in Nestorowa and Paul data. The following table is the genes and their altered magnitude. For the full list of leukemia related genes, see Extended Data Table 1 in [30] and Supplemental Table S4 in [31].

| Up-regulated<br>Gene | log2-fold<br>change | Down-regulated<br>Gene | log2-fold<br>change |
| --- | --- | --- | --- |
| CD34 | 2.1500 | LGALS3 | -3.4901 |
| LAPTM4B | 1.8000 | CYBB | -2.9546 |
| MMRN1 | 1.3600 | CD36 | -2.7661 |
| SOCS2 | 1.2400 | ANXA5 | -2.6349 |
| CDK6 | 1.2300 | LY86 | -2.5564 |
| CPXM1 | 1.2000 | IRF8 | -2.4982 |
| EMP1 | 1.0100 | SAMHD1 | -2.4580 |
| GPR56 | 2.7004 | GRN | -2.3659 |
| GATA2 | 1.8875 | RNASE6 | -2.3585 |
| LPIN1 | 1.6323 | FCER1G | -2.2934 |
| MZB1 | 1.4854 | S100A9 | -2.2447 |
| ZSCAN18 | 1.3219 | TLR4 | -2.1078 |
| GUCY1A3 | 1.2630 | FCGRT | -2.1016 |
| SPNS2 | 1.2016 | S100A8 | -2.0116 |
| PTK7 | 1.2016 | CLEC12A | -1.8730 |
| ABCC1 | 1.1375 | MNDA | -1.8417 |
| SYTL1 | 1.0704 | IL13RA1 | -1.7515 |
| MAGED1 | 1.0704 | SGK1 | -1.7418 |
| ARHGAP25 | 1.0704 |  |  |
| SLA2 | 1.0000 |  |  |

**S4 Fig.**
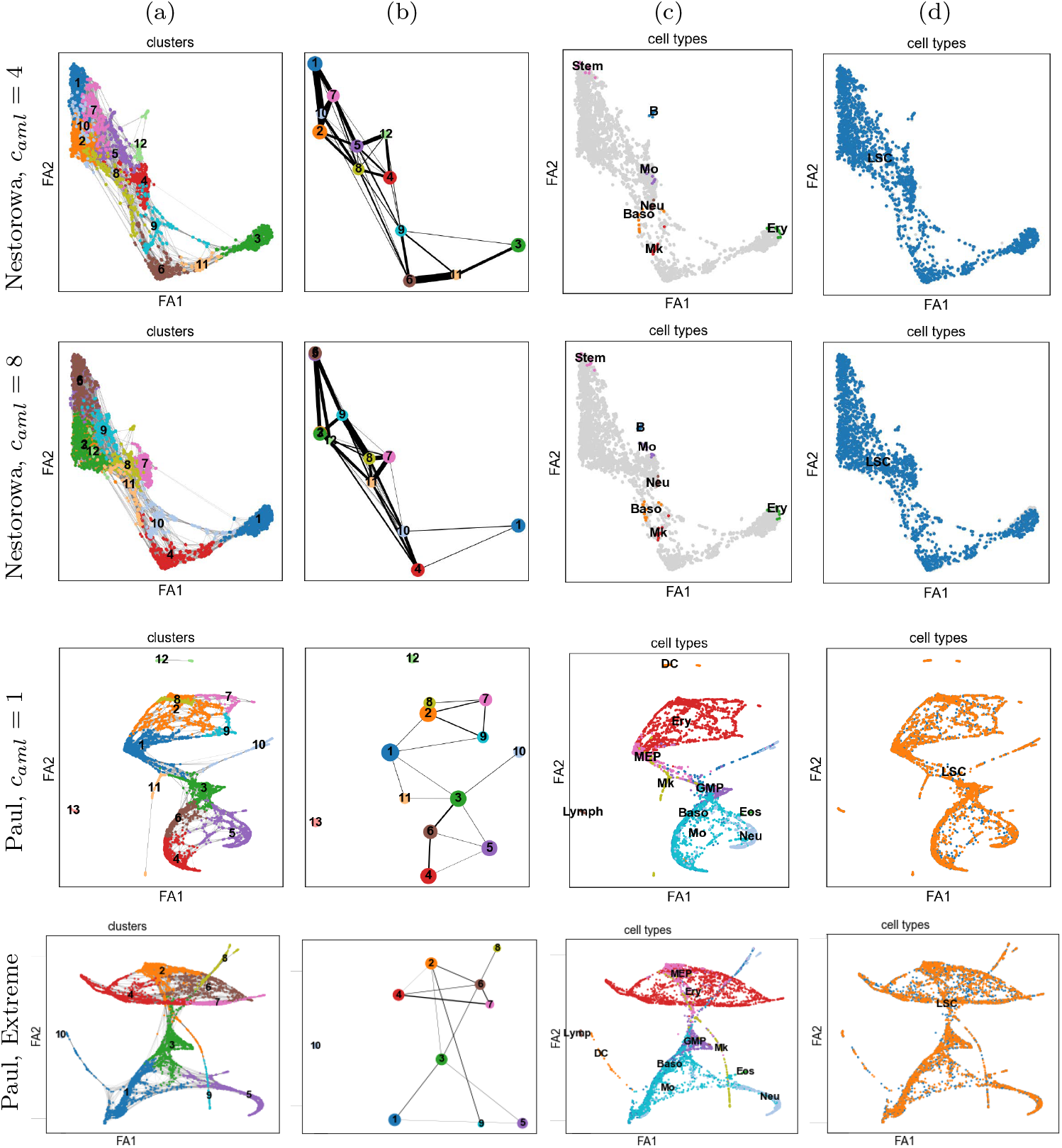
Graph computed with altered genes in larger magnitude. The graphs are computed from leukemic genes altered with larger magnitudes *c*_*r*_ = 4 and 8, where 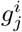 is modified as 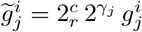. From the regular leukemic alteration *γ*_*j*_, we consider additional alterations of *c*_*r*_. The shown figures are the clusters (a), abstracted graph (b), annotated normal cells (c), and leukemic cells (d), comparable to Fig 7. The PAGA algorithm does not distinguish the AML cells to the regular cells despite the larger magnitudes of change in Nestorowa data. We observe that in Paul data, the graph algorithm does not distinguish even the extreme level of altered AML cells.

**S5 Fig.**
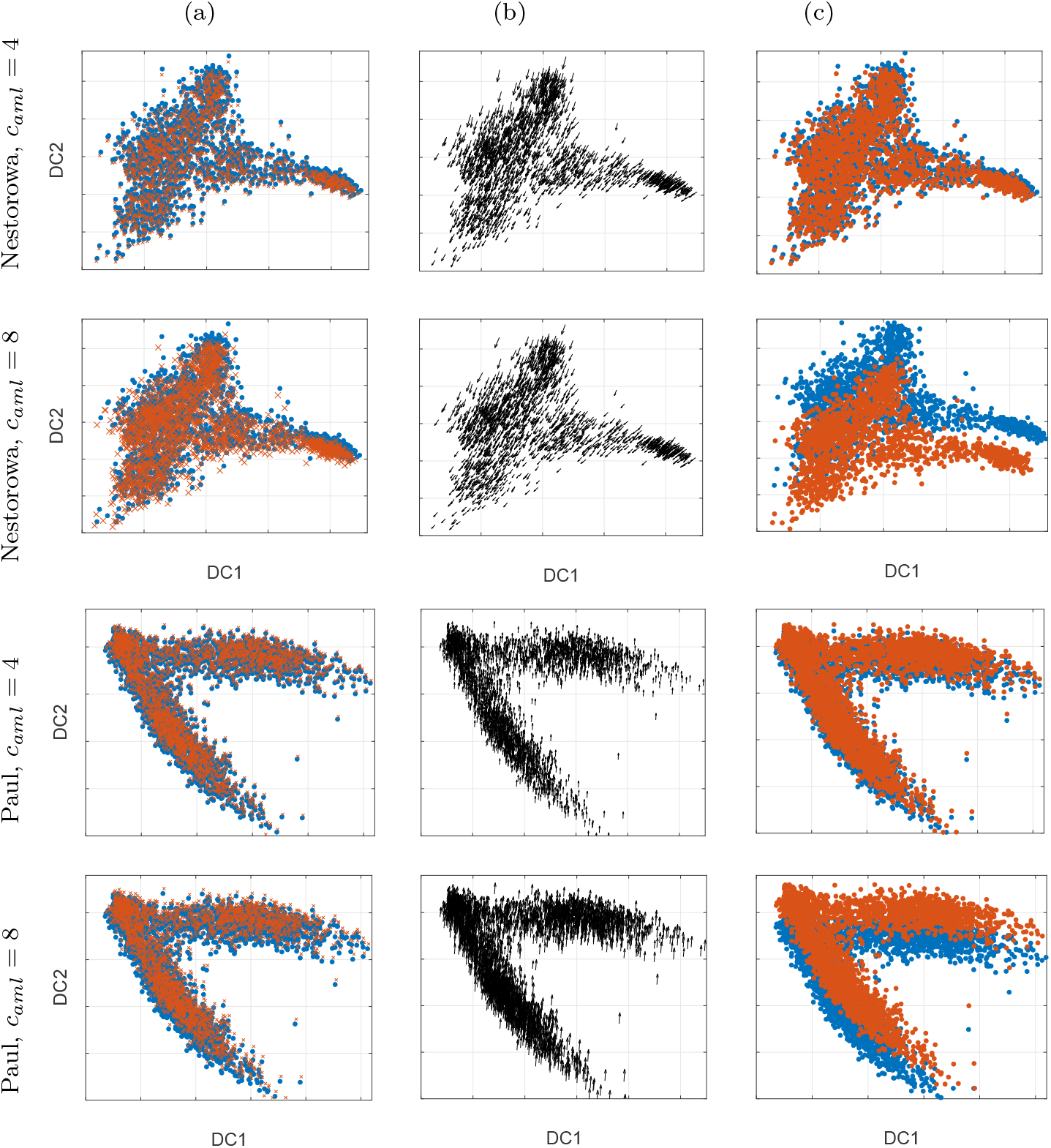
Reduced component space computed with altered genes in larger magnitude. Similar to S4 Fig, the following diffusion component spaces and direction of leukemia pathogenesis are computed from leukemic genes altered with larger magnitudes *c*_*r*_ = 4 and 8. The shown figures are projection of perturbed single cell (x) on the normal diffusion component space (a), and its altered direction toward stem cells (b). The reduced diffusion space with both normal and perturbed cells (c) show more apparent difference in the projected altered cells and also in the reduced component space, however in a similar direction.

**S6 Fig.**
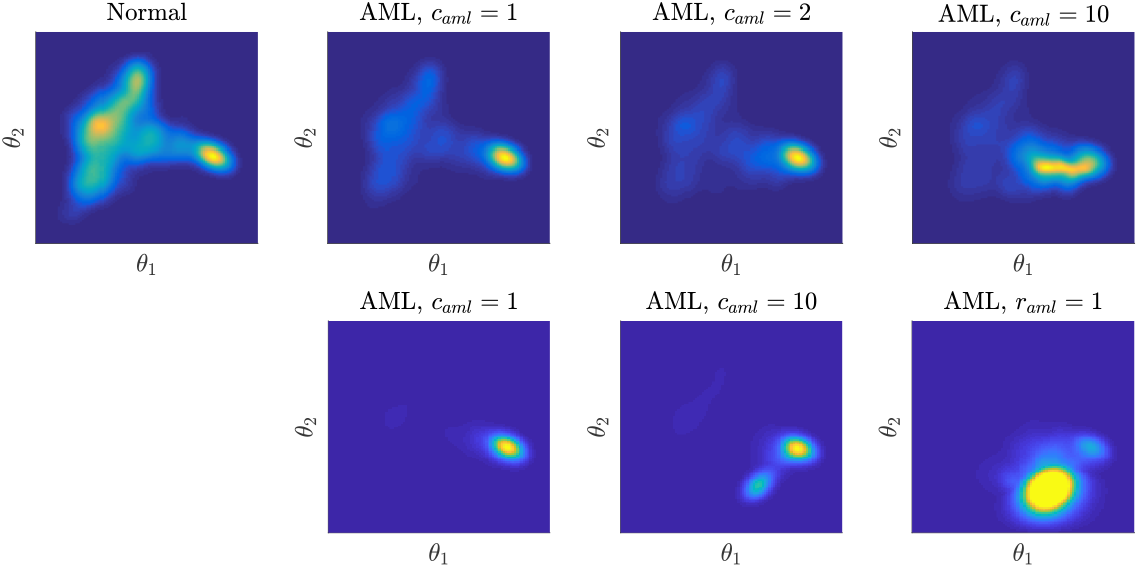
Abnormal cell landscape during leukemia pathogenesis. The distribution of cell states *u*(*t, θ*) at *t* = 20 show abnormal cell states emerging during leukemic progression. The presented figures are computed by using different advection models as 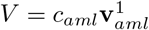 (top) and 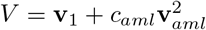 (bottom), with various levels of *c*_*aml*_ = 1, 2, 10. Larger magnitude of *c*_*aml*_ results in more disrupted cell landscape.

**S7 Fig.**
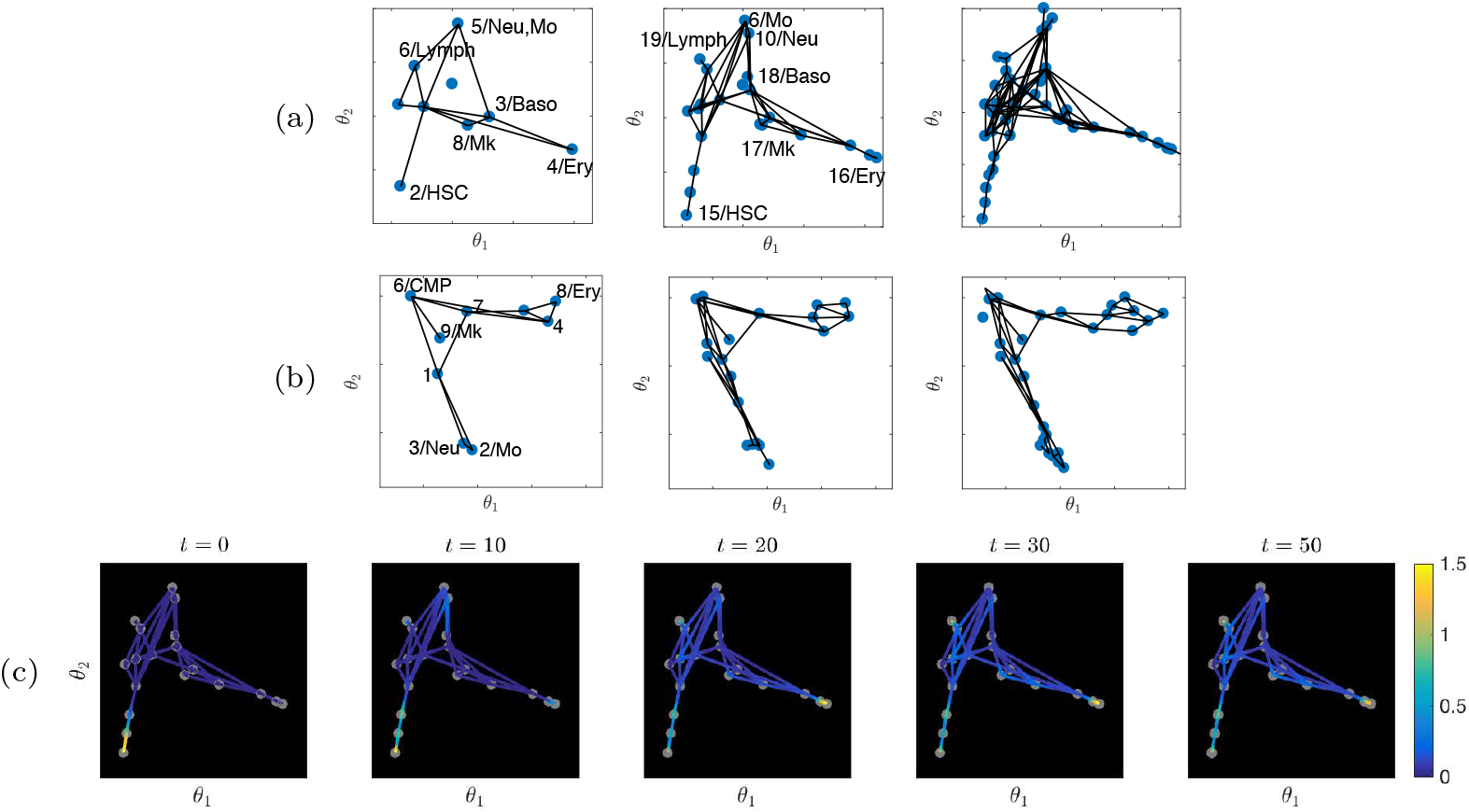
Graph refinement. The hierarchy of graphs using partition-based graph abstraction [17] and single cell data from Nestorowa et al. (2016) (a) and Paul et al. (2015) (b). The single-cell data can be regarded as the most refined graph. The simulation of normal hematopoiesis on graph with 19 nodes (c) is comparable to Fig 3(a).

**S8 Fig.**
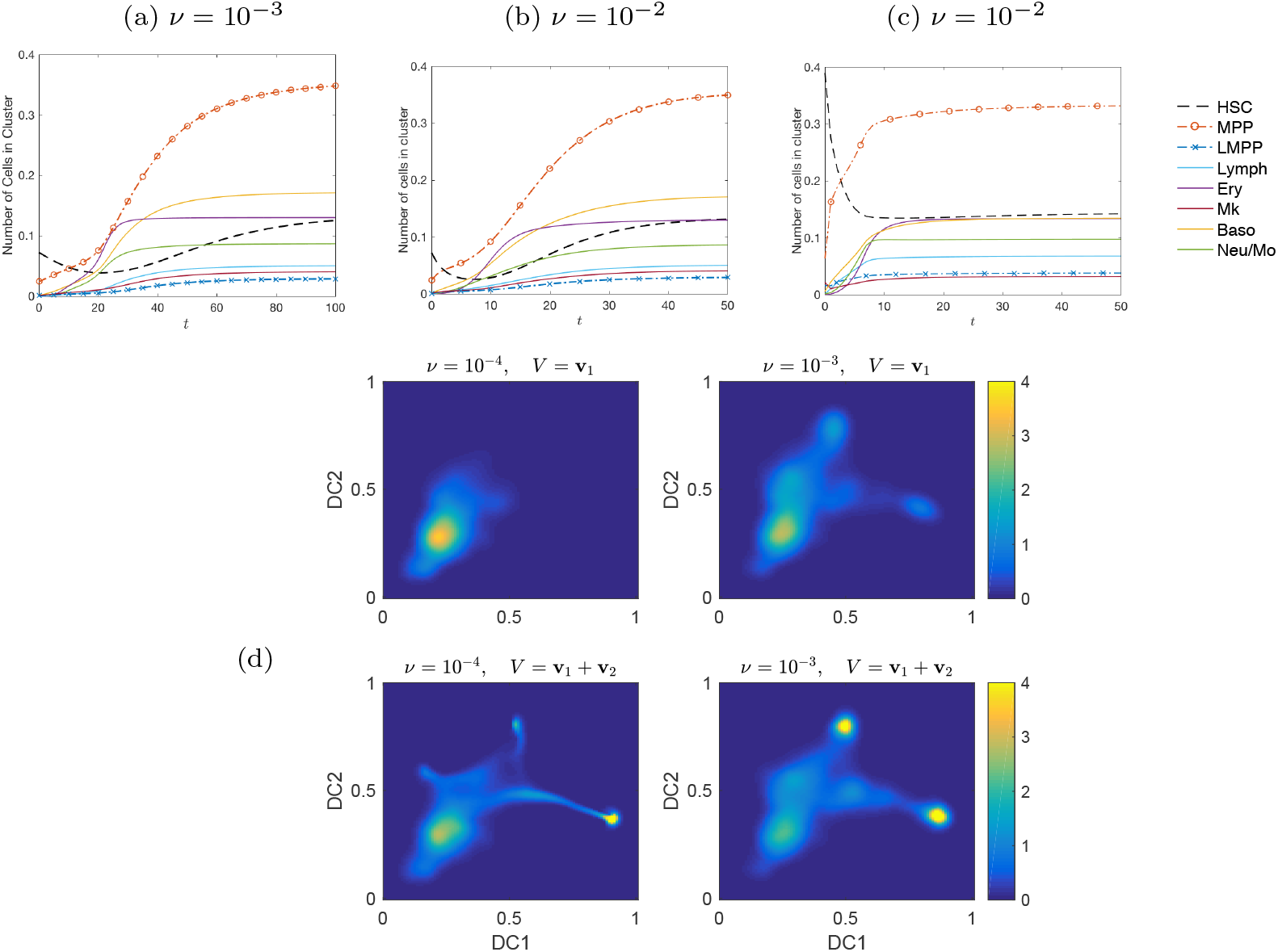
Model sensitivity to parameters. Using single cell data from Nestorowa et al. (2016), number of cells and its dynamics in each cluster up to *t* = 50 for different values of *ν* and initial stem cell numbers *ρ*(0) are shown in (a–c). The dynamics of cells in each cluster for *ν* = 10^*−*3^ with *ρ*(0) = 0.1 (a), *ν* = 10^*−*2^ with *ρ*(0) = 0.1 (b), and *ν* = 10^*−*2^ with larger initial number of cells, *ρ*(0) = 0.5 (c) shows that the recovery is more rapid for larger values of *ν* and larger number of initial stem cells. Cell distribution *u*(*θ, t*) at intermediate time *t* = 14 for advection terms **v**_1_ and **v**_1_ + **v**_2_, and *ν* = 10^*−*4^ or *ν* = 10^*−*3^ are shown in (d). Larger values of *ν* increases overall rate of differentiation, while adding **v**_2_ prioritizes recovery of the most matured cells.

## Acknowledgments

Research reported in this publication was supported by the National Cancer Institute of the National Institutes of Health under award number P30CA033572. The content is solely the responsibility of the authors and does not necessarily represent the official views of the National Institutes of Health.

